# Integration and comparison of multi-omics profiles of NGLY1 deficiency plasma and cellular models to identify clinically relevant molecular phenotypes

**DOI:** 10.1101/2021.05.28.446235

**Authors:** Songjie Chen, Guangwen Wang, Xiaotao Shen, Daniel Hornburg, Shannon Rego, Rene Hoffman, Stephanie Nevins, Xun Cheng, Michael Snyder

## Abstract

NGLY1 (N-glycanase 1) deficiency is a rare congenital recessive disorder of protein deglycosylation unaddressed by the current standard of care. Using combined metabolomics and proteomics profiling, we show that NGLY1 deficiency activates the immune response and disturbs lipid metabolism, biogenic amine synthesis, and glutathione metabolism. These alterations were also observed in NGLY1 deficient patient-derived induced pluripotent stem cells (iPSCs) and differentiated neural progenitor cells (NPCs), which serve as personalized cellular models of the disease. These findings provide molecular insight into the pathophysiology of NGLY1 deficiency and suggest potential therapeutic strategies.

## Introduction

NGLY1 deficiency is a rare inherited recessive disorder which gives rise to complex symptoms, including global developmental delay, delayed bone maturation, movement disorders, alacrimia, peripheral neuropathy, hepatopathy, and seizures^1–6^. Symptoms arise in infancy and vary between individual patients. Most patients do not survive past childhood^7, 8^ and no therapeutic options have been developed thus far. As revealed by whole exome sequencing, the disorder is caused by a mutation in *NGLY1* gene which encodes the cytosolic enzyme N-glycanase1 (NGLY1) responsible for protein deglycosylation, a process necessary for the endoplasmic reticulum associated degradation (ERAD) of misfolded or damaged glycoproteins^1, 9–12^. At the molecular level, N-linked glycosylation directs the proper folding of glycoproteins during their insertion into the endoplasmic reticulum (ER) ^10, 13, 14^. It also plays an essential role in diverse protein functions, including the targeting of glycoproteins to appropriate destinations, regulating their stability, solubility, and antigenicity, and modulating receptor binding kinetics^15–18^. The inactivation of NGLY1 disrupts the homeostatic degradation of glycoproteins and associated modification of transcription factors, giving rise to the diverse NGLY1 disorders^16, 19, 20^.

Recent studies of the disorder have discovered several key substrates of NGLY1, including the nuclear factor erythroid 2-related factor 1 (NRF1) and bone morphogenic protein 4 (BMP4) ^16, 17, 20–22^. NRF1 is reported to regulate proteasomal activity^23^, inflammatory responses, lipid homeostasis^24, 25^ and diverse metabolic processes^26, 27^. BMP4 is involved in the differentiation of various critical cell types, such as neurons ^28–31^.

To further clarify the affected lipidome, metabolome, proteome, and immunome alterations induced by NGLY1 deficiency, we used a multi-omics approach to characterize the dysregulation of downstream pathways and identify potential therapeutic targets. Specifically, we applied lipidomics and targeted metabolomics profiling of patient plasma to characterize NRF1-mediated modulation of lipid metabolism; untargeted metabolomics profiling to identify additional metabolism alterations; and proteomics profiling to characterize NGLY1 deficiency-relevant proteins, affected signaling pathways, and immunome dysregulation. By integrating multi-omics profiles, we found molecular phenotypes with the potential to provide detailed mechanistic insights and guide therapeutic development and clinical translation.

However, the heterogeneity of NGLY1 syndromes, its rarity, and the lack of prior study has limited our understanding of and therapeutic options for NGLY1 deficiency. Thus, we also generated induced pluripotent stem cells (iPSCs) from a subset of patients and applied CRISPR to generate isogenic controls. The patient-derived iPSCs enhanced the exploration of underlying mechanisms and provided detailed molecular insight. As NGLY1 deficiency patients demonstrate multiple neural specific symptoms, we further differentiated these iPSCs into neural progenitor cells (NPCs) to trace the progression of the disorder in an *in vitro* model of the nervous system^32–34^. This panel of undifferentiated and differentiated iPSCs were then profiled using both targeted and untargeted metabolomics and proteomics. By integrating multi-omics profiles of plasma and iPSCs/NPCs samples at the molecular level, we established a new research paradigm for the study of the underlying pathology mechanisms of NGLY1 deficiency and identified several pathways for further research and testing of therapeutic strategies.

## Results

### Summary of cohort and research design

An overview of the study is shown in Fig.1. With the support of the Grace Science Foundation (https://gracescience.org/), Stanford University generated a cataloged biobank of NGLY1 patient families’ biological samples, genomic information, and medical records (https://med.stanford.edu/news/all-news/2017/10/biobank-entrepreneur-accelerate-research-into-rare-disease.html). This study was conducted with 58 plasma samples from the biobank, covering 19 diagnosed NGLY1-deficient patients and 39 healthy immediate family members (Fig. 1A, B, Table 1). To the best of our knowledge, this is the most detailed and largest NGLY1 deficiency cohort available to date. In this study, we performed comprehensive molecular profiling of metabolites (metabolome), lipids (lipidome), proteins (proteome), and immunoproteins (immunome) of plasma collected from patients and healthy controls.

**Fig 1.**
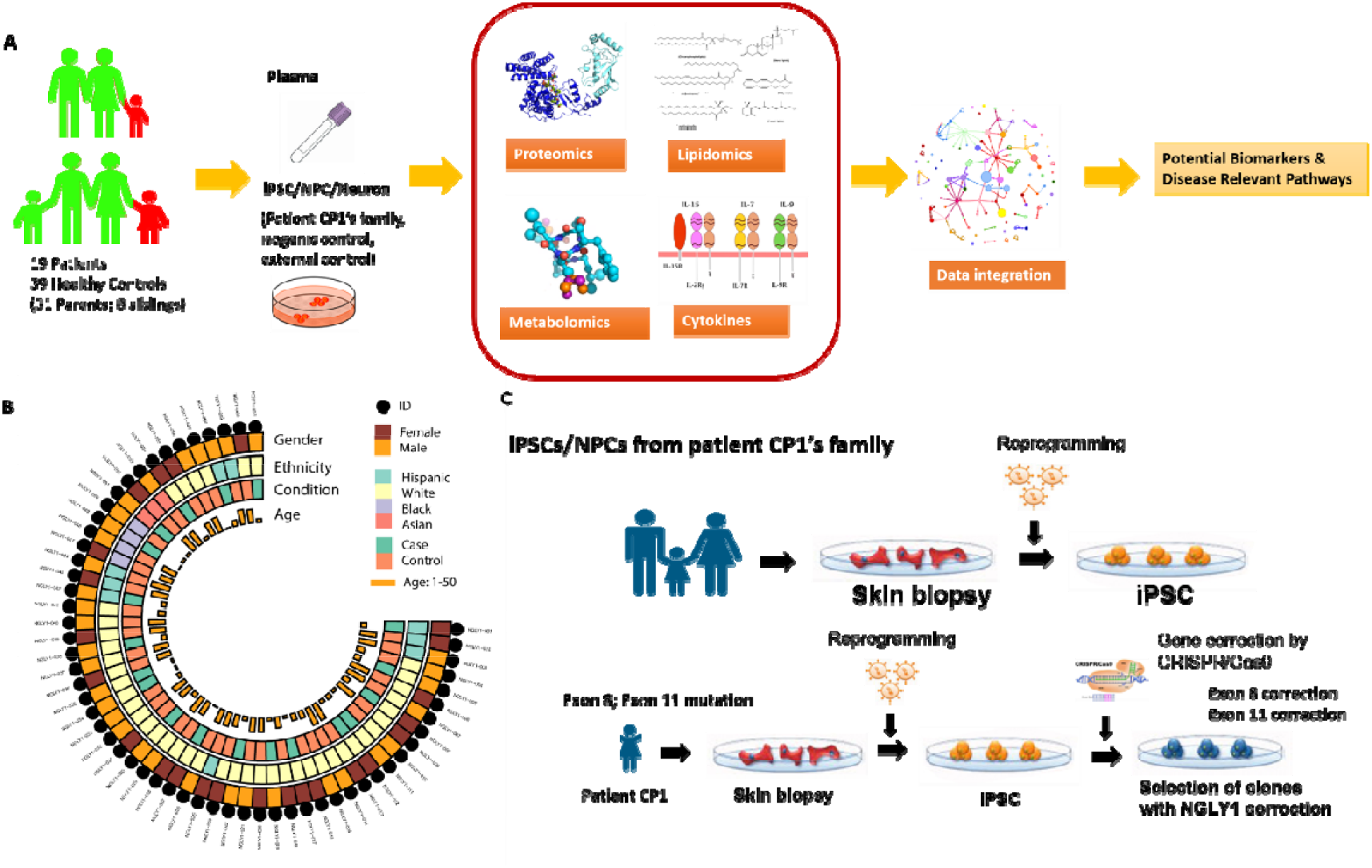
Overview of the multi-omics profiling process. A) Multi-omics profiling of NGLY1 deficiency cohort and patient-derived cellular models. A unique cohort including 19 NGLY1 deficiency patients and 39 of their healthy family members were recruited for metabolomics and proteomics analysis. We collected plasma from the whole cohort and derived iPSCs from patient CP1’s family. The plasma proteome, metabolome, lipidome, and cytokines were quantified to identify potential therapeutic targets for NGLY1 deficiency. We also profiled the metabolome and proteome of patient CP1’s iPSCs/NPCs and their isogenic and external controls to validate the alternation of multi-omics profiles during the development of NGLY1 deficiency. B) The demographic characteristics of the patient cohort, showing patients and controls’ gender, age and ethnicity. C) Overview of the personalized cellular disease models developed from patient CP1’s family samples and modified by CRISPR technology.

**Table 1.**
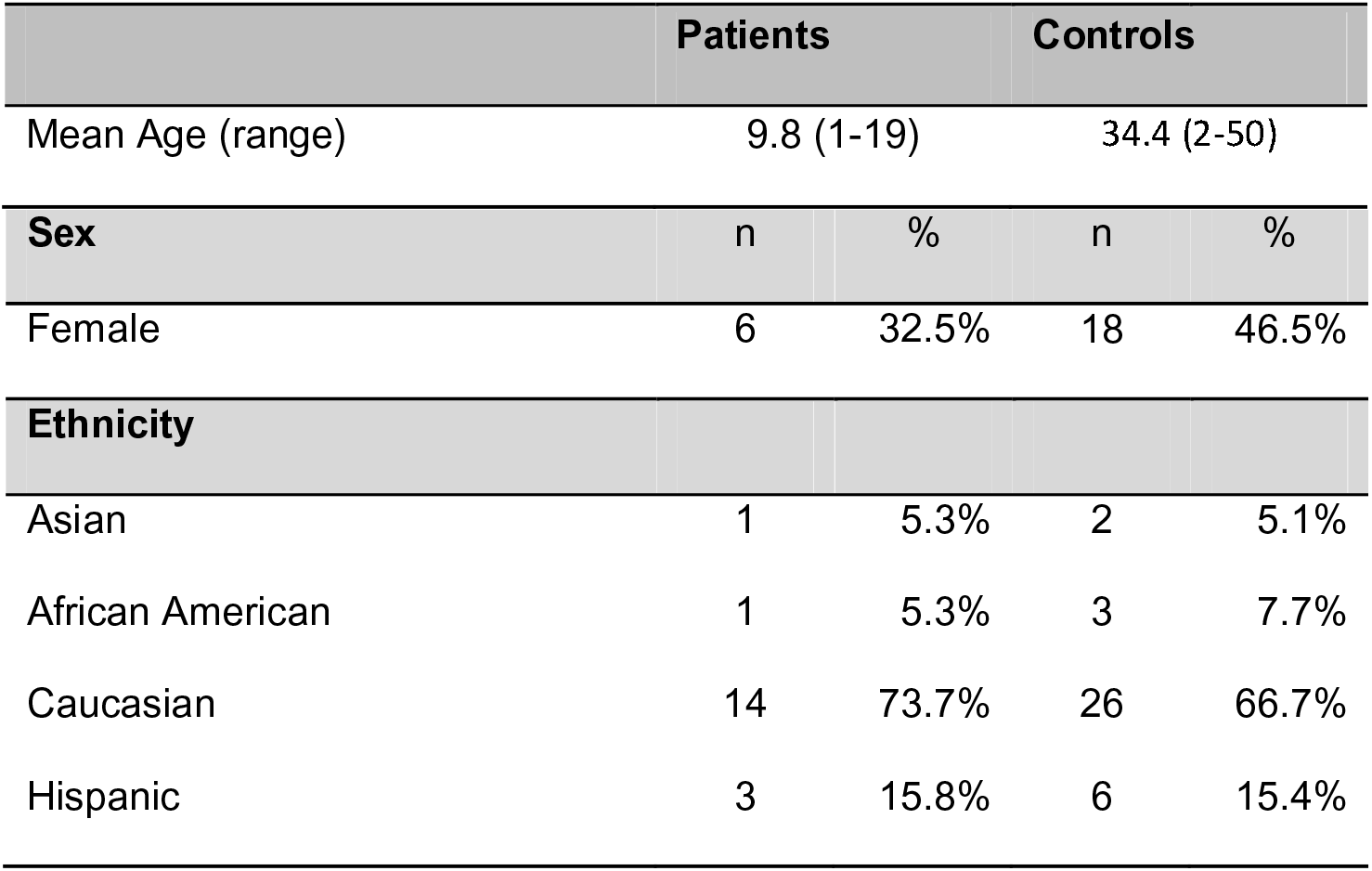
Characteristics of the NGLY1 deficiency cohort. Summary of plasma samples collected in the NGLY1 deficiency biobank. Patient gender, age, and ethnicity were further adjusted by mixed linear model.

To further characterize disease-associated metabolites, proteins, and pathways, we generated a series of iPSCs from fibroblasts of patient CP1’s family (CP1, CP1Fa, CP1Mo) and an external healthy control (MSJ) and then differentiated the iPSCs into NPCs via dual inhibition of Smad signaling (Fig. 1C, 2, Table 2, Fig. 2) ^35^. CP1 carries two unique mutations of *NGLY1* in each allele (exon 8: p.402_403del; exon 11: p.R524X). The patient-derived iPSCs were characterized by their expression of pluripotency markers (Supplementary Fig. 1A) and karyotyping (Supplementary Fig. 1B). Preservation of mutations in *NGLY1* in CP1’s iPSCs was confirmed by Sanger sequencing (Fig. 2A). We also leveraged CRISPR/Cas9 technology to edit the genotype of the cells at individual CP1 alleles, enabling evaluation of whether correction in either mutation (CP1-E8C; CP1-E11C) or both mutations (CP1-E118C; CP1-E811C) is necessary and sufficient to rescue the molecular phenotype in the iPSC models. Recovery of the *NGLY1* gene in CP1’s iPSCs was confirmed by Sanger sequencing (Fig. 2B). Patient plasma metabolome, lipidome, proteome (Supplementary Table 2-8), cellular metabolome, and proteome data were integrated to identify and define molecular phenotypes associated with NGLY1 deficiency.

**Fig 2.**
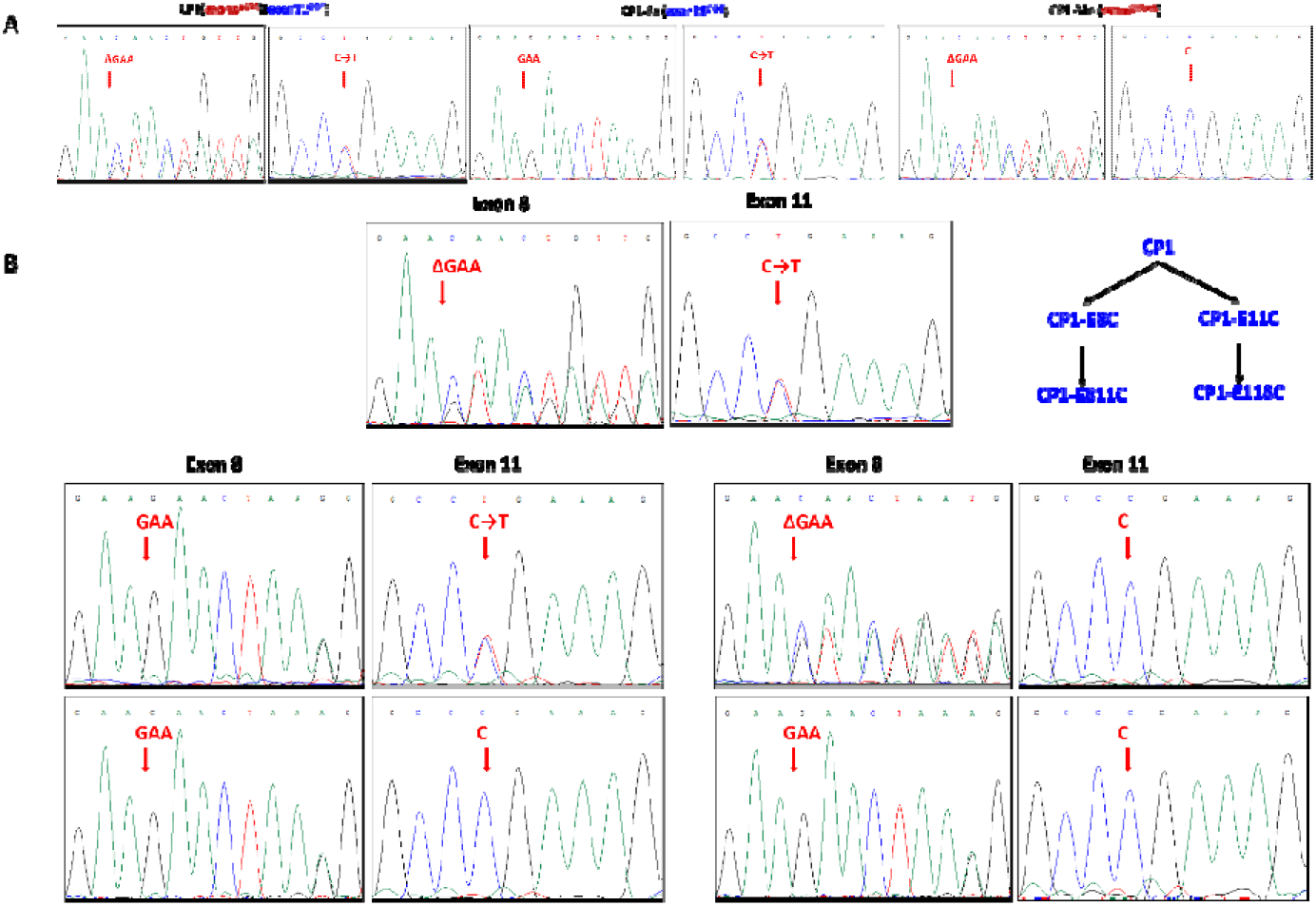
Characterization of patient-derived iPSCs. A) Confirming NGLY1 mutations in iPSCs by Sanger sequencing. B) Confirming the correction of NGLY1 mutations by CRISPR in iPSCs.

**Table 2.**
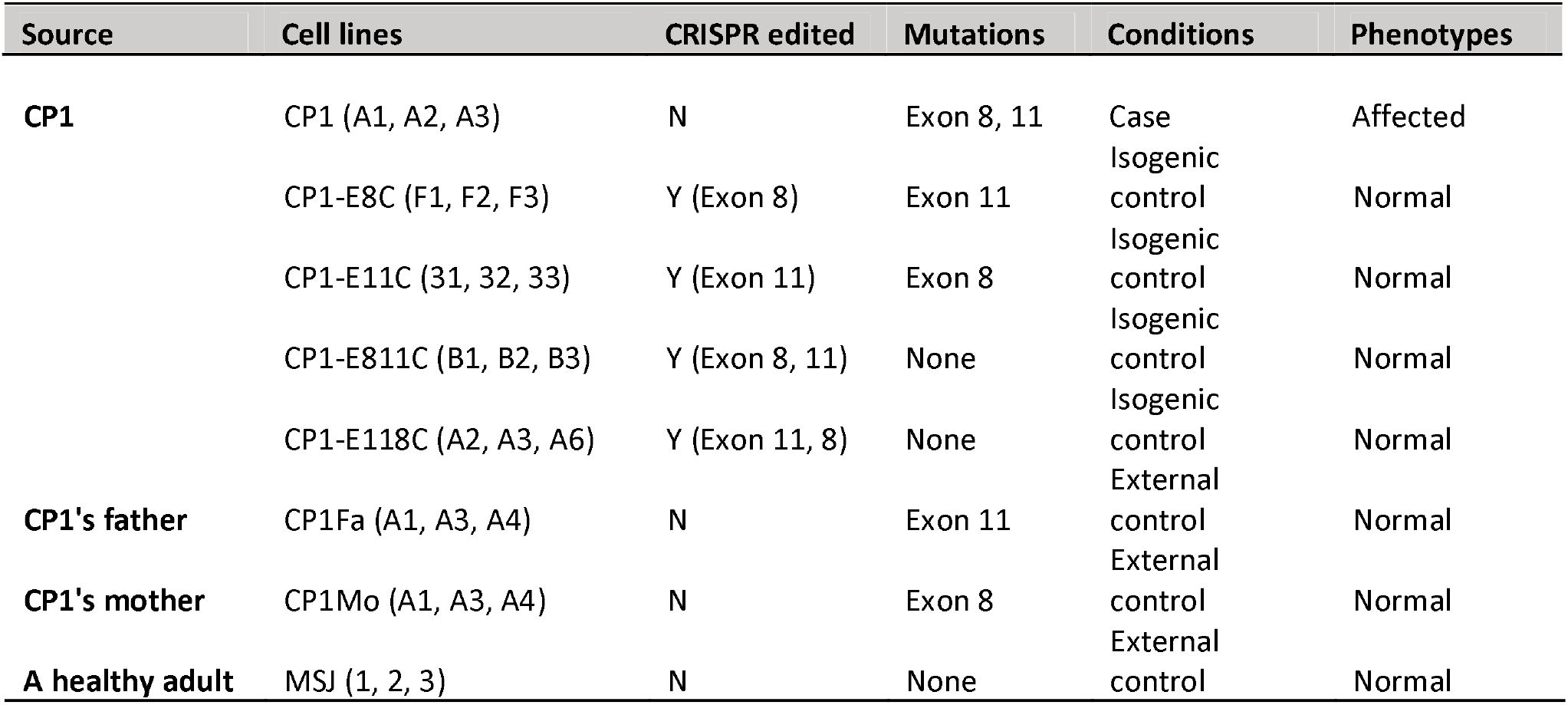
iPSC/NPC generated from patient CP1’s family. Summary of iPSCs/NPCs generated from patient CP1’s family. Each cell line includes three clones (n = 3). Patient CP1 carries two alleles with mutations in exon 8 and 11. Four CRISPR-corrected CP1 cell lines (CP1-E8C, CP1-E11C, CP1-E811C, CP1-E118C) served as isogenic controls. Cell models derived from patient’s healthy parents and an unrelated healthy adult were used as external controls.

### Targeted metabolomics and lipidomics quantified NGLY1 deficiency induced dysregulation of lipid metabolism

#### Overview of targeted metabolomics and lipidomics profiles suggest global downregulation of lipids in NGLY1 patients’ plasma

We conducted deep profiling of the NGLY1 cohort’s plasma metabolome and lipidome to characterize alterations of lipid metabolism induced by NGLY1 deficiency. 396 metabolites and lipids were quantified using the Absolute IDQ-p400 kit (Biocrates Life Science AG) ^36^ (Supplementary Table 3). 723 lipids were quantified by Lypidyzer (Sciex) (Supplementary Table 5). Based on the targeted metabolomics profiles and lipidomics profiles, hierarchical clustering (Supplementary Fig. 3A, 4A) and principal component analysis (PCA) (Supplementary Fig. 3B, 4B) demonstrated partial separation between patients and controls. As shown in the volcano plot of the metabolome and lipidome (Supplementary Fig. 3C, 4C), most lipids were downregulated in NGLY1 patients’ plasma.

To further adjust for potential bias induced by differential age distributions between patients and controls (Table 1, Fig. 1B), a mixed linear regression model was applied to quantified metabolites. A total of 69 quantified metabolites demonstrated significant correlation with NGLY1 deficiency (P < 0.05) (Supplementary Fig. 3D, Supplementary Table 3). These metabolites were classified as amino acids, biogenic amines, glycerophospholipids, acylcarnitines (AC), sphingolipids (SP), cholesterol esters (CE), diacylglycerols (DAG), and triacylglycerols (TAG). The same regression model also identified 90 lipids quantified by lipidomics with significant correlation to the disorder (P < 0.05), including CE, DAG, TAG, ceramide (CER), dihydroceramide (DCE), free fatty acids (FFA), lysophosphatidylcholines (LPC), lysophosphatidylethanolamine (LPE), phosphatidylcholines (PC) AND phosphatidylethanolamines (PE), sphingomyelin (SM) (Supplementary Fig. 4A, 4B, 4D, Supplementary Table 5).

#### Functions of dysregulated lipids showed correlations with symptoms

The targeted metabolomics and lipidomics profiling of NGLY1 plasma samples showed consistent downregulation across a broad panel of plasma lipids, including LPC, PC, CE, CER, SM, DAG, TAG, etc., except for PC 40.6, which was slightly elevated in NGLY1 patients’ plasma (Supplementary Fig 3C, 4B).

PCs are the key component of the lipid bilayer of cells and are involved in diverse metabolism and signaling pathways^37^. Although most PCs ere downregulated in patients’ plasma, the function of elevated PC 40.6 level in NGLY1 deficiency-associated lipid metabolism is unknown. CEs are a major component of blood lipoprotein particles used for lipid transport and storage ^38–40^. A recent study suggested potential correlation between cardiovascular disease and inefficient synthesis of CE from cholesterol^41^. Abnormal cholesterol metabolism is a typical phenotype of a disrupted nuclear factor erythroid 2-related factor 2 (NRF2) pathway^42^, which is involved in the activation of DNA damage responses and could be a compensatory mechanism induced by NGLY1 deficiency and NRF1 dysregulation^43^. In particular, the downregulated plasma CEs in NGLY1 patients are mainly odd-chain fatty acids (CE 17.2, CE 19.2, CE 19.3, Supplementary Fig. 3D), indicating a potential dysregulation in fatty acid synthesis and lipid transport. LPCs are components of oxidized low-density lipoproteins, usually involved in atherosclerosis and inflammatory diseases^44, 45^. The downregulated LPCs in NGLY1 patients’ plasma suggest low overall inflammation (Supplementary Fig. 3C, 3D, 4A, 4D), thus, the immune response of NGLY1 deficiency becomes interesting. Several SMs in patients’ plasma were also downregulated across diverse fatty acid chain lengths. SMs are highly enriched in plasma membranes and involved in signal transduction^46, 47^. SM-CER metabolism changes have been previously associated with diverse diseases, such as neurodegenerative diseases, cardiovascular diseases, and cancers^48–50^. CER species were also slightly downregulated in NGLY1 patients’ plasma (Supplementary Fig 4A, 4D). Both SM and CER belong to the broader SP class, which is crucial for the biosynthesis of glycosphingolipids and gangliosides. Notably, the fatty acid chains of the downregulated lipids exhibited a wide range of chain lengths and saturation, although most free fatty acids did not show significant variation between NGLY1 patients and controls. Overall, fatty acid metabolism was generally downregulated in NGLY1 patients’ plasma, which suggests potential mitochondrial dysfunction.

#### Elevated acylcarnitines and relevant proteomics alternations suggest potential interruption of lipid transport

Several ACs were upregulated in patients’ plasma (AC 2.0, AC 14.0, AC 16.0, etc. Fig. 3A, Supplementary Fig. 3A), which indicated dysregulation of mitochondrial fatty acid metabolism and activation of related stress responses^51^. In NGLY1 patients’ plasma proteome, a paralog of the formylglycine-generating enzyme (SUMF2, Fig 3A) was significantly upregulated, which is involved in the metabolism of SPs and their interactions with interleukins^52, 53^. Cytoplasmic aspartate aminotransferase (GOT1) and transketolase (TKT) were also elevated in NGLY1 patient’s plasma (Fig 3A), indicating increased aspartate aminotransferase activity in hepatic neoglucogenesis and adipocyte neoglycerogenesis^54–56^. From this, the transport processes of myriad lipids or lipid precursors may be disrupted by NGLY1 deficiency. As most patients are reported to exhibit abnormal liver functions^1, 57^, GOT1 and TKT could serve as potential targets for mitigating hepatic symptoms. The downregulation of cholic acid in patients’ plasma indicates reduced fat absorption and cholesterol excretion^58, 59^. This metabolic change may also affect the dysregulation of other carbohydrate metabolism pathways, which is supported by elevated levels of lactate and deoxyribose 5-phosphate as well as reduced levels of glycerol-3-phosphate (Supplementary Fig 5B-D). In addition, the reduced overall lipid metabolism, combined with the elevated activity of glycolysis intermediates, indicate potential global mitochondrial dysfunction^60^, which may affect systemic energy metabolism of NGLY1 patients (Fig 3B-D).

**Fig 3.**
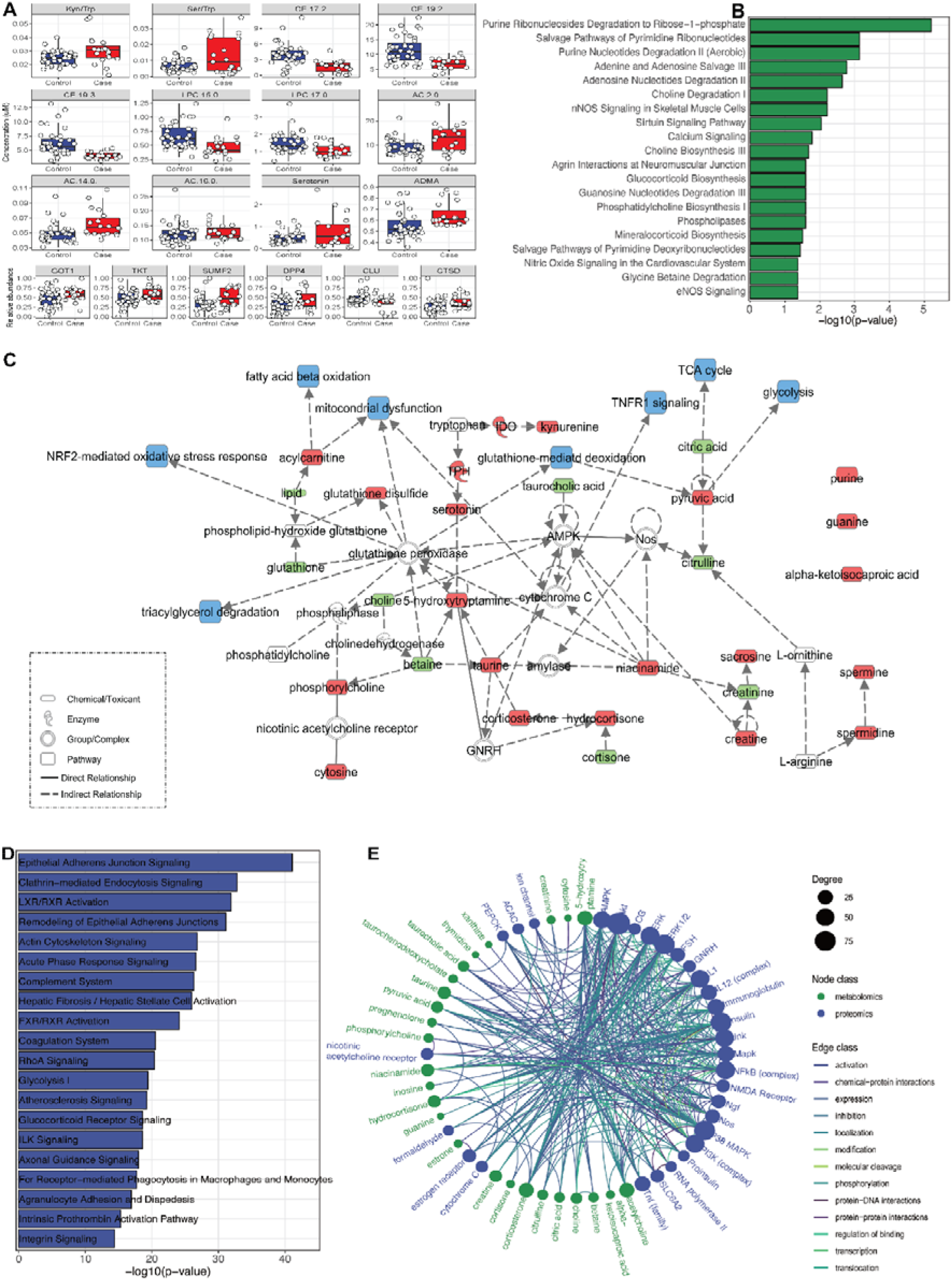
NGLY1 deficiency-induced metabolome and proteome dysregulation in plasma. A) Selected plasma metabolites and proteins with significant variation between controls (blue) and patients (red) (P < 0.05). B) Enrichment of dysregulated pathways in the plasma metabolome (top 20). C) Diagram of dysregulated plasma metabolites involved pathways. The upregulated metabolites/proteins are shown in red; the down regulated metabolites/proteins are shown in green; affected pathways and functions are shown in blue. D) Enrichment of dysregulated pathways of the plasma proteome (top 20). E) Networks composed of dysregulated proteins and metabolites in patients’ plasma. Nodes re metabolites and proteins with significant alteration in patient plasma. The edges show the direct and indirect relations between nodes.

### Untrageted metabolomics and proteomics profiling characterized more pathway alternations

#### Overview of untargeted metabolomics profiles of plasma and patient-derived cells

We further profiled a broad spectrum of NGLY1 deficiency-relevant plasma metabolites using untargeted mass spectrometry and annotated 1091 putative metabolic features using an in-house library and a combined public spectral database (https://doi.org/10.1101/2021.05.08.443258), from which 76 metabolic features were annotated at a metabolomics standards initiative (MSI) level 1 or 2^61^. Although hierarchical clustering and PCA only demonstrated minor separation between NGLY1 patients and controls (Supplementary Fig. 5A, 5B), a panel of metabolites with level 1 or 2 identification showed significant variation (P < 0.05) between controls and patients (Supplementary Fig. 5D). For the profiling of disease-modeling iPSCs, we captured 101973 metabolic features with putative annotation. In iPSCs, 181 metabolites were annotated with level 1 or 2 identification while in NPCs, 89 metabolites were annotated with level 1 or 2 identification. Based on the untargeted metabolomes of NGLY1 iPSCs and NPCs, the hierarchical clustering and PCA showed a relatively clear separation between the patient’s cells, isogenic control cells, and healthy adult cells.

#### Overview of proteomics profiles of plasma and NGLY1 patient-derived iPSCs/NPCs

In addition to metabolites and lipids, we also used a standardized TMT labeling protocol to quantify NGLY1 patients’ plasma proteins^62, 63^. Although hierarchical clustering and PCA of the plasma proteome was not able to significantly separate patients and controls (Supplementary Fig. 7A, 7B), a broad panel of plasma proteins demonstrated significant variation and fold change between patients and controls (Supplementary Fig. 7C, 7D), including diverse immune-relevant proteins. In cellular proteomics profiles, we quantified 3449 proteins in iPSC lysates and 4940 proteins in NPC lysates (Supplementary Table 12, 13). However, the hierarchical clustering and PCA of iPSC and NPC proteomes did not show clear separation between patients and controls (Supplementary Fig. 8A-C, 9A-C), Supplementary Fig. 8D and 9D show iPSC and NPC proteins with most significant fold changes.

#### NGLY1 deficiency patients show dysregulated creatine and amino acid metabolism

Compared with the targeted plasma profiles, untargeted plasma profiling found similar alterations in amino acid and biogenic amine derivative levels. Creatinine, as a biomarker of glomerular filtration rate, was significantly downregulated in NGLY1 patients’ plasma (Supplementary Fig. 3A, 3C, 3D). Its upstream and downstream metabolites, phosphocreatine and sarcosine, respectively, were slightly upregulated (Supplementary Fig. 3A, 3D, 5B, 5D), suggesting an enhanced turnover rate of creatine metabolism. However, abnormal kidney function has not been previously reported in NGLY1 patients and only one case with adrenal insufficiency has been seen^64^, suggesting these effects may be modest.

Diverse amino acids (valine, phenylalanine, tryptophan, asparagine, lysine, threonine, proline) and their derivatives, such as N-omega-hydroxy arginine and pyroglutamic acid, were slightly downregulated in patients’ plasma, whereas several polyamines including putrescine, spermidine, and sarcosine—which can act as growth factors for cell division— were upregulated in NGLY1 patients’ plasma (Supplementary Fig. 3A, 3C, Fig. 3C). Tryptophan metabolism plays a critical role in the modulation of immune responses as well as the regulation of growth and neurodegeneration^65, 66^. Although we did not observe significant alternation of tryptophan derivatives, the ratio of kynurenine/tryptophan was significantly upregulated in patients’ plasma (Fig. 3A), suggesting elevated activity of indoleamine-2,3-dioxygenase (IDO), an enzyme that plays a role in a variety of pathophysiological processes such as antimicrobial and antitumor defense, neuropathology, immunoregulation, and antioxidant activity. The kynurenine pathway also modulates carbohydrate metabolism^67–69^. The ratio of serotonin/tryptophan was also upregulated in patients’ plasma (Fig. 3A), suggesting increased activity of the tryptophan hydroxylase (TPH), which catalyzes the first step of serotonin synthesis. Serotonin is an important hormone and neurotransmitter whose dysregulation impacts diverse functions in gut-brain signaling, memory, sleep, and behavior. The elevation of serotonin level varies among NGLY1 patients (Fig. 3A) which may associate with the heterogeneity of neural syndromes. This pathway is associated with an elevated risk for a variety of neurological disorders, including schizophrenia, and somatic anxiety^70, 71^.

#### NGLY1 deficiency patient show dysregulated redox activity

NGLY1-related redox dysregulation was also characterized in untargeted metabolomics profiling, which revealed slight downregulation of glutathione and S-adenosylmethionine balanced against slight upregulation of cysteinylglycine disulfide, pyroglutamic acid, and asymmetric dimethylarginine (ADMA) (Fig 3A, Supplementary Fig. 3A, 5B, 5C). Asymmetric dimethylarginine (ADMA) is a metabolic by-product of protein modification processes found in the cytoplasm of all human cells. It interferes with arginine in the production of nitric oxide, which is a key signal involved in endothelial and cardiovascular health^72^. The methyl groups transferred to create ADMA are derived from the methyl group donor S-adenosylmethionine, an intermediate in the metabolism of homocysteine and polyamines^73, 74^ (Fig. 3C, Supplementary Fig. 3A, 5B-D). Remarkably, several proteins detected in plasma proteome, including GOT1, TKT, SUMF2, DPP4, CLU and CTSD, also play a role in the above pathways, showing a systemic metabolic response to NGLY1 deficiency (Fig. 3A, 3C). Among these proteins, CLU prevents aggregation of stress-induced protein aggregations^75–77^; DPP4 is also involved in the activation of T-cell and NF-κB^78–81^; CTSD is required for amyloid precursor protein degradation and associated with Alzheimer’s disease^82^.

#### Pathway analysis of NGLY1 plasma metabolome and proteome data suggests potential therapeutic strategies

To further investigate NGLY1 deficiency-induced phenotypes, pathway enrichment analysis was performed on plasma metabolome and proteome data. The metabolomic pathways significantly affected by NGLY1 deficiency were mainly involved in purine nucleotide degradation, skeletal muscle cell signaling, choline degradation, and phospholipase activity (Fig. 3B). Compared with the altered metabolomics pathways, a greater number of proteomic pathways were affected by NGLY1 deficiency, including epithelial adherens junction signaling, endocytosis signaling, LXR/RXR activation, actin cytoskeleton signaling, and acute phase response signaling, etc. (Fig 3D). The top enriched pathways in the plasma proteome and metabolome profiles overlapped with tissue organization, lipid metabolism, and immune signaling. In the plasma proteome, adherens junction signaling plays an essential role in the development of normal tissue organization^83^. LXRs are important regulators of cholesterol, fatty acid, and glucose homeostasis, which may be involved in the lipid transport and glucose generation changes noted above^84, 85^. Actin cytoskeleton signaling is crucial for movement regulation and influences NF-κ phase response plays an essential role in the innate immune system, cross-talking with LXR signaling in response to inflammatory cytokine signaling. By merging the top enriched pathways of the plasma proteome with those seen in the metabolome, we obtained a comprehensive network of NGLY1-related changes comprising neurotransmitters, lipid metabolites, cytoskeleton proteins, and cytokines with diverse relationships (Fig. 3C, 3E). Several complexes and enzymes demonstrated essential roles in the merged network, such as AMP-activated protein kinase (AMPK), cytochrome C, and glutathione peroxidase, which may serve as potential therapeutic targets.

#### iPSC/NPC metabolome and proteome data show similar pathway alterations as plasma data

The personalized iPSC/NPC models derived from patient CP1’s family were used as a model to study the underlying disease pathology. The NGLY1 iPSC metabolome and proteome demonstrated several significant differences between patients and controls that affected diverse pathways, including nucleotide degradation, amino acid metabolism, EIF2 signaling, protein ubiquitination pathways, epithelial adherens junction signaling pathways, and myriad signaling pathways (Fig. 5A).

**Fig 4.**
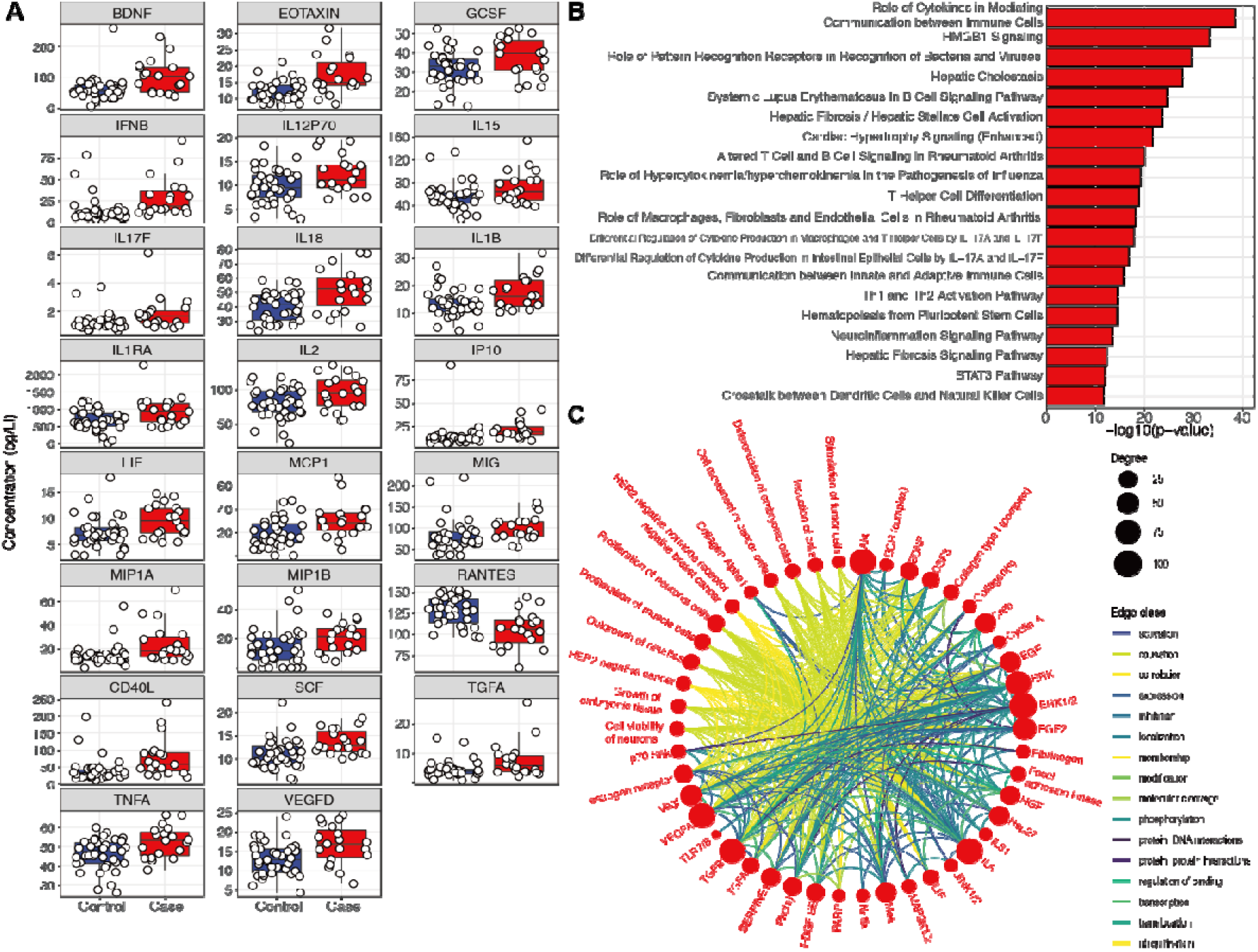
NGLY1 deficiency induced dysregulation of the cytokine pathway. A) Cytokines with most significant fold changes and variation between controls (blue) and patients (red). B) Enrichment of dysregulated cytokine pathways. The bar shows -log_10_ P value. C) Summary of the most upregulated cytokines and their relevant functions and diseases in the network. The edges show the direct and indirect relationships between each two cytokines.

**Fig 5.**
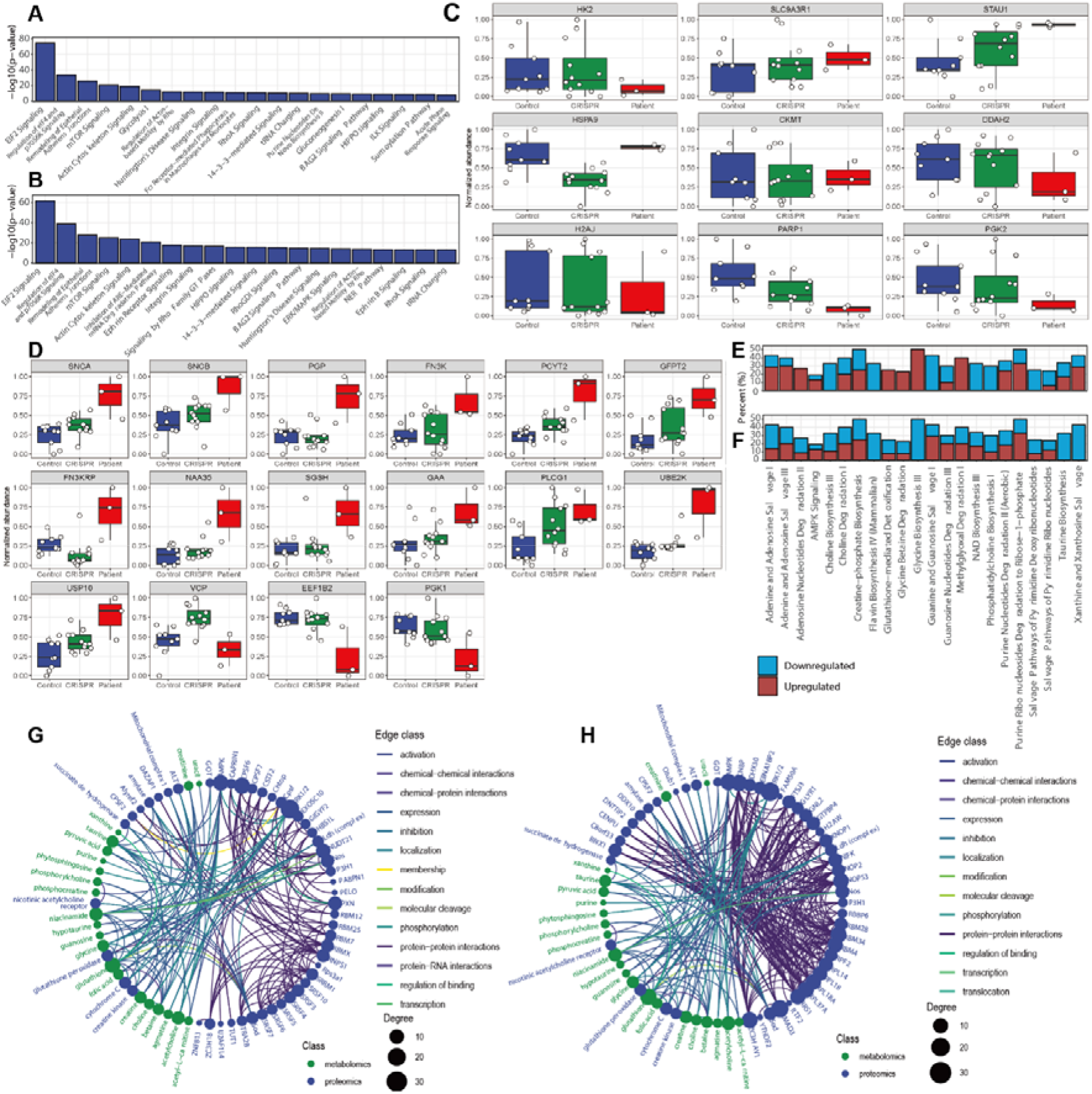
NGLY1 deficiency pathway dysregulation in patient-derived iPSCs/NPCs. A) Dysregulated proteomics pathways in patient-derived iPSC (-log_10_ P > 5). The bar shows the P value. B) Dysregulated proteomics pathways in patient-derived NPC (-log_10_ P > 5). The bar shows the -log_10_ P value. C) iPSC proteins with significant variation between patient (blue), CRISPR-corrected isogenic control (green) and external control (red). D) NPC proteins with significant variation between external control, CRISPR-corrected isogenic control and patient. E) Dysregulated metabolomics pathways in patient-derived iPSC (P < 0.05). The bar shows the percentage of downregulated and upregulated metabolites in the pathway. F) Dysregulated metabolomics pathways in patient-derived NPC (P < 0.05). The bar shows the percentage of downregulated and upregulated metabolites in the pathway. G) Networks comprising dysregulated proteins (blue) and metabolites (green) in patient-derived iPSC. Nodes are metabolites and proteins with significant alterations. The edges show the direct and indirect relations between each node. H) Summary of the networks comprising dysregulated proteins and metabolites in patient-derived NPCs. Nodes re metabolites and proteins with significant alterations. The edges show the direct and indirect relationships between each node.

The iPSC proteome also included elevation of stress response and immune signaling proteins. The enrichment of EIF2 signaling pathway incorporates different stress-related signals to regulate alternative mRNA translation^88, 89^. Other pathways affected in NGLY1 iPSCs included the epithelial adherens junctions and actin cytoskeleton signaling pathways and were also dysregulated in NGLY1 plasma proteome. Several enriched iPSC pathways were directly linked with protein degradation, such as the unfolded protein response, which was upregulated, whereas protein kinase A signaling was downregulated. Several metabolic pathways, such as glycolysis and purine nucleotide synthesis, also overlapped with the affected pathways in the plasma metabolome.

The metabolomic profiles of iPSCs were consistent with the patient plasma metabolomes, with most amino acids, amino acid derivatives, and lipids, especially free fatty acids, downregulated in patient iPSCs. The modulation of nucleotide metabolism is more complex, with downregulation of guanine and guanosine balanced against upregulation of adenine, adenosine, and hypoxanthine (Supplementary Fig. 6A, 6C).

NPC proteomes showed a greater number of significant pathway alterations between NGLY1 patients and controls. Interestingly, the most significantly dysregulated pathways in iPSCs were also dysregulated in NPCs, including EIF2 signaling, protein ubiquitination pathways, and epithelial adherens junction signaling pathways. (Fig. 5B). Other relevant pathways included proteasome-mediated degradation pathways, protein ubiquitination, interleukin signaling, cellular response to stress, and diverse signaling pathways. In addition to the signaling and immune relevant pathways which partially overlapped with the pathways identified in iPSCs and plasma (Fig. 5A, 3D, 4B), the NPC proteome demonstrated greater alternation of pathways relevant to protein degradation (Fig. 5B). Unlike iPSC proteome, most of the quantified NPC proteins were upregulated in patient’s NPCs (Fig. 5D, Supplementary. 9D).

The NPC metabolome also showed significant downregulation of amino acids and derivatives, indicating a globally down-regulated protein metabolism. Comparison of the NPC metabolome with iPSC metabolome showed that the top enriched pathways were identical; although the upregulated and downregulated metabolites were nonetheless distinct (Fig 5E, 5F). In the NPC metabolome, greater downregulation of a broader array of metabolites involved in xanthine and xanthosine salvage, taurine biosynthesis, and adenine nucleotide degradation pathways was seen whereas creatine-phosphate biosynthesis and choline degradation pathways displayed similar levels as in NGLY1 plasma and iPSC.

#### Proteins identified in iPSCs/NPCs function in plasma pathways

Among the iPSC proteins with the most significant variation between NGLY1 patients and controls (Fig. 5C), the majority were ribosomal proteins, scaffold proteins, and heat shock proteins (SLC9A3R, STAU1, HSPA9). Several enzymes, such as HK2, DDAH2, PARP1, and PGK2, were downregulated in patient iPSCs. HK2 is a hexokinase that phosphorylates glucose to produce glucose-6-phosphate, the first step in glucose metabolism. The downregulation of H2K may also interact with the dysfunction of mitochondria and dysregulation of lipid metabolism^90^. DDAH2, as a dimethylarginine dimethylaminohydrolase, regulates the cellular concentrations of methylarginines and generates nitric oxide. It is associated with chronic kidney disease and preeclampsia^91, 92^. PARP1 (poly(ADP-ribosyl)transferase) is a chromatin-associated enzyme that modifies various nuclear proteins and DNA and is involved in the regulation of various important cellular processes such as differentiation, proliferation, and antiviral innate immunity^93^. PGK2 (Phosphoglycerate kinase 2) is phosphoglycerate kinase that catalyzes the reversible conversion of 1,3-bisphosphoglycerate to 3-phosphoglycerate, broadly affecting carbohydrate metabolism^94^. H2AJ is a replication-independent histone that is a variant H2A histone, which inhibits diverse signaling pathways such as Wnt signaling^95^. SLC9A3R1 is a scaffold protein that connects plasma membrane proteins and regulates their surface expression^96^. CKMT, which is a mitochondrial creatine kinase responsible for the transfer of high energy phosphate from mitochondria creatine, was also slightly upregulated in patient’s iPSCs. Thus, we observed upregulation of phosphocreatine in patient iPSCs^97^ (PMID: 26708229), which could explain the reduced plasma creatinine levels previously observed.

In the NPC proteome, synuclein alpha (SNCA) and synuclein beta (SNCB) were both upregulated in NGLY1 patient NPCs (Fig. 5D). SNCA, as a biomarker for neurodegenerative diseases, is abundant in the brain with smaller amounts found in the heart, muscles, and other tissues. Although the function of SNCA is not well understood, studies suggest that it plays a key role in maintaining an adequate supply of synaptic vesicles in presynaptic terminals^98, 99^. SNCA peptides are a major component of amyloid plaques in the brains of patients with Alzheimer’s disease. Its aggregation is affected by PC composition in cell membranes, whose components were also altered by NGLY1 deficiency^100, 101^. SNCB is a non-amyloid component of senile plaques found in Alzheimer disease. It may also act as a regulator of the SNCA aggregation process^102, 103^. Although no direct observation of protein aggregation in NGLY1 deficiency cells has been previously reported, SNCA-associated synaptic vesicle trafficking and cytoskeleton deficiency may account for observed features in the neurological phenotype of NGLY1 deficiency.

NPC proteomes demonstrated dysregulation across a broad spectrum of ubiquitinylation-relevant proteins (Fig 5D), although some of their roles in NGLY1 deficiency remain uncharacterized. For example, UBE2K as a ubiquitin-conjugating enzyme that mediates the selective degradation of short-lived and misfolded proteins. It also determines neurogenic potential during prenatal and postnatal brain development^104^. Upregulated USP10 was also observed, which belongs to a ubiquitin-specific protease family of cysteine proteases. The enzyme specifically cleaves ubiquitin from ubiquitin-conjugated protein substrates and can thus activate AMPK^105, 106^. A previous study suggested a connection between ubiquitinylation, AMPK activity, and NGLY1 deficiency^107^. The reduced VCP is an ATPase that emerge a central role in the ubiquitin system, involved in protein degradation, DNA repair and replication, regulation of the cell cycle, activation of the NF-κB pathway, and also causes strong mitochondrial uncoupling^108, 109^.

A panel of enzymes involved in carbohydrate and lipid metabolism were also altered in NGLY1 patient NPCs (Fig. 5D). PCYT2, for instance, catalyzes the formation of CDP-ethanolamine from CTP and phosphoethanolamine in the Kennedy pathway of phospholipid synthesis^110^. GFPT2 (glutamine-fructose-6-phosphate transaminase 2) controls the flux of glucose in the hexosamine pathway. It may be involved in regulating the availability of precursors for N- and O-linked glycosylation of proteins^111^. Similar to the NGLY1 iPSC proteome, phosphoglycerate kinase PGK1 was also downregulated in patient NPCs^112^. Another down-regulated protein in the patient NPCs, EEF1B2, acts as a translation elongation factor in the transfer of aminoacylated tRNAs to the ribosome. It is associated with diverse diseases and neurodevelopmental disorders^113^.

Among the dysregulated cytosol enzymes, N-Alpha-Acetyltransferase 35 (NAA35) is auxiliary component of the N-terminal acetyltransferase C (NatC) complex which catalyzes N-terminal acetylation, a process important for the maintenance of protein function and cell viability^114, 115^. The elevated Fructosamine 3 Kinase (FN3K) and Fructosamine 3 Kinase Related Protein (FN3KRP) catalyze the phosphorylation of fructosamines of glycated proteins, further resulting in deglycation^116^. They are also modulators of NRF2 activity and are involved in the response to oxidative stress by mediating deglycation of NRF2^117^. PLCG1 catalyzes the formation of inositol 1,4,5-trisphosphate and diacylglycerol from phosphatidylinositol 4,5-bisphosphate, contributing to cell death and inflammatory reactions^118, 119^. GAA is an alpha glucosidase, which is essential for the degradation of glycogen to glucose in lysosomes. Its deficiency leads to the glycogen accumulation in lysosomes, also known as Pompe’s disease, which is an autosomal recessive disorder characterized by muscle weakness similar to NGLY1 deficiency^120, 121^. Related pathways include innate immune system activation and glucose metabolism, which are also affected by NGLY1 deficiency. SGSH is a sulfamidase involved in the lysosomal degradation of heparan sulfate. This dysfunction is associated with lysosomal storage disorders resulting from impaired degradation of heparan sulfate, which is relevant to early onset neurodegeneration^122^.

### Cytokine profiles demonstrate elevated activity of the immune system

To further characterize the NGLY1 deficiency-induced immune response, we conducted targeted quantification of 62 plasma cytokines. 23 cytokines demonstrated significant variation between patients and controls by linear mixed regression (P < 0.05) (Supplementary Table 8, Supplementary Fig 8C). However, hierarchical clustering and PCA of cytokine profiles did not show a clear separation between patients and controls (Supplementary Fig 8A, 8B). Unlike the plasma lipidome, immune-related proteins were upregulated in patient plasma, indicating a global activation of immune responses (Fig. 4A). The most significant upregulated cytokine was BDNF, which plays an important role in the regulation of the neuroimmune axis and is associated with schizophrenia, Alzheimer’s disease, mood disorders, and Parkinson’s disease^123, 124^. Eotaxin was also significantly elevated in patient plasma. This cytokine is related to allergic reactions and Th2-skewed immune response (Fig. 4A, Supplementary Fig. 8). Eotaxin shapes the differentiation of neural progenitor cells and microglia, which are involved in the progression of aging, neurogenesis, and neurodegeneration; the latter more commonly associated with NGLY1 deficiency. Increased circulating levels of eotaxin have been observed in major psychiatric disorders (schizophrenia, bipolar disorder, major depression) ^125^. Other upregulated cytokines included interleukins, leukemia inhibitory factor, chemokines, and interferons, which are involved in chemotactic activation of inflammatory and NF-κB pathways^126^. In contrast, RANTES was downregulated in NGLY1 patient plasma. RANTES mediates monocyte/macrophage infiltration into atherosclerotic lesions^127^. Previous studies also suggested that dysregulation of the immune system is closely associated with broad alterations in lipid metabolism^128^.

Cytokine profiles suggested dysregulation of communication between immune cells, as seen in HMGB1 signaling and hepatic cholestasis with elevated levels of most cytokines in the network (Fig. 4B, C). These disrupted pathways impair diverse cellular functions, such as the cell viability of neurons, differentiation of embryonic cells, and growth of the brain. The canonical NF-κB complex was also activated by both cytokines and other plasma proteins such as DPP4 and may perform an essential role in the NGLY1 affected cytokine network (Fig. 4C).

### Integration of multi-omics profiles identified molecular phenotypes associated with syndromes

By comparing the enrichment of metabolomic and proteomic profiles across different sample types, we found more similar pathway regulation patterns between NGLY1 patient plasma and patient-derived NPCs (Fig. 6A, 6B). The metabolic pathways relevant to cell proliferation, lipid metabolism, amino acid metabolism, and a panel of signaling pathways showed similar modulation in all three samples. Compared with iPSCs, NPCs demonstrated a more similar regulation pattern to patient plasma for pathways involved in immune response, response to oxidative stress, and signaling pathways affecting lipid metabolism and neurodegeneration. These findings suggest an overarching switch of pathway regulation during the progression of NGLY1 deficiency. Common pathways with similar regulation behavior in both iPSCs and NPCs were mainly associated with nucleotide synthesis and cell viability (Fig. 6A, 6B), suggesting potential proliferation and viability deficiency in NGLY1 cases. We further mapped the dysregulated proteins identified in plasma, iPSCs, and NPCs to a tissue-specific proteome database^129^ (Fig. 6C) The enrichment of NGLY1-affected proteins in different tissues suggested a correlation with tissue-specific syndromes, such as movement disorders, hepatopathy, peripheral neuropathy, multifocal epileptiform activity and cerebral atrophy. We found that NGLY1-associated proteins were enriched in the brain, liver, muscle and heart^57, 64, 131, 132^. Although most of these enriched proteins only show minor alternations in patient plasma and cell models, the enrichment of these proteins in specific tissues could induce pathological syndromes. The iPSC/NPC metabolome and proteome also add more information of key intermediates and enzymes of the plasma metabolic pathways, such as the upregulation of glutathione peroxidase and CKMT, and the involvement of AMPK and cytochrome C (Fig. 6D), providing a potential for therapeutic intervention. A graphic summary of the molecular phenotypes identified by metabolomics and proteomics profiling of different sample types and associated tissue-specific syndromes is shown in Fig 6E, which provides more insights into potential mechanisms of NGLY1 induced syndromes. Besides the common metabolomics and proteomics phenotypes associated with nervous, hepatic and muscular syndromes, specific molecular phenotypes relevant to immune response, neurodegeneration and kidney dysfunction are also discovered. However, the direct mediator between NGLY1 deficiency and the molecular phenotypes remains unclarified.

**Fig 6.**
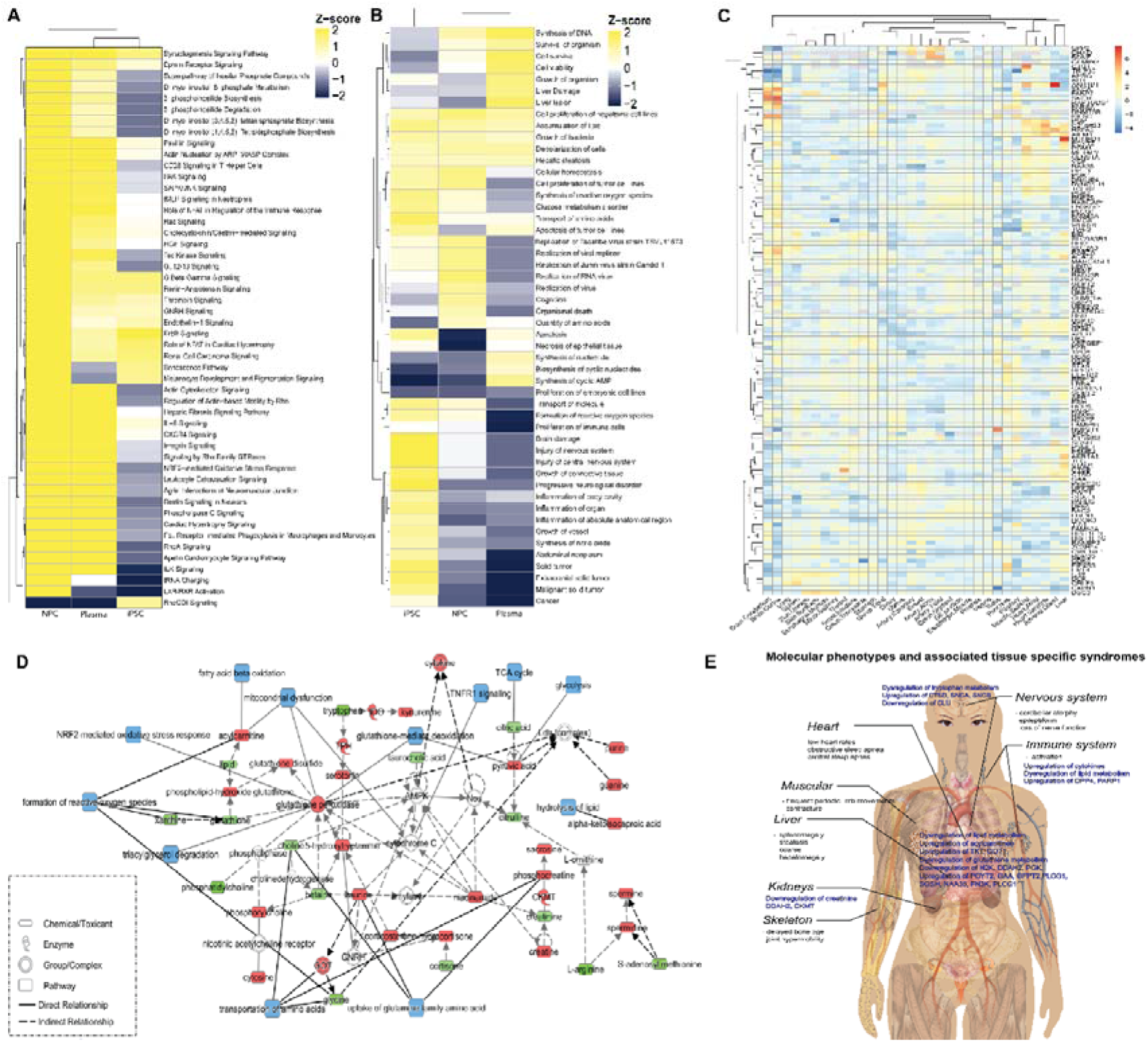
The summary of NGLY1 deficiency induced pathway disruption. A) The clustering of disrupted functional metabolome pathways in patients’ plasma and patient-derived iPSC and NPC. B) The clustering of disrupted proteome pathways in patients’ plasma and patient- derived iPSC and NPC. C) Tissue enrichment of NGLY1 associated proteins identified in patients’ plasma and patient-derived iPSC and NPC. D) NGLY1 affected metabolic pathways annotated by integrating metabolome and proteome data of plasma, iPSC and NPC. The upregulated metabolites/proteins are shown in red; the down regulated metabolites/proteins are shown in green; affected pathways are shown in blue. E) A graph summary of identified molecular phenotypes and associated syndromes.

## Discussion

The integration of multi-omics profiles remains a technical challenge despite the rapid development of complementary data analysis strategies^133, 134^. Unlike population data-driven investigations, the study of rare diseases is constrained by small patient cohort sizes. Thus, patient-derived personalized cellular models provide a complementary model to address the sample size limitation. The isogenic controls generated by CRISPR technology added even more power for multi-omics profiling and data analysis. We observed minor differentiation between the external control and isogenic control in iPSCs/NPCs (Fig. 5C, 5D, Supplementary Fig.6 A-D, Fig 8 A, 8B, Fig. 9A, B). No significant differentiation was observed between CRISPR corrected heterozygote and homozygote iPSCs/NPCs. The differentiation of iPSCs to NPCs enables the characterization of myriad pathway alterations during disease development.

Although the NGLY1 deficiency-associated downregulation of amino acid derivatives was a common phenotype across both patient plasma and cellular models, the plasma phenotype was less significant after applying linear regression to adjust for age and gender. Nonetheless, clinical records indicated that most patients took multivitamin and amino acid dietary supplements. Thus, even if masked by patient plasma supplementation, the downregulation of amino acid metabolism could be a more significant phenotype, with the potential to interrupt protein degradation or cell proliferation. The downregulation of lipids was also a common phenotype across the different types of samples; however, the affected lipids varied substantially and the overall downregulation of lipids was less significant in the cellular models. In this, the global downregulation of the plasma lipidome may be a function of interrupted molecular trafficking. Fatty acids were the most common down-regulated lipid species, although the chain length and saturation in different sample types varied significantly. Taking into account of the fatty acid chains across diverse lipid species, NGLY1 deficiency appeared to induce a general and global dysregulation of fatty acid synthesis, which could be due to disrupted carbohydrate metabolism that shares intermediates with fatty acid synthesis or the interruption of fatty acid metabolism and transport.

Biogenic amine synthesis is a complicated network with extensive crosstalk between different super-pathways. Glutathione was downregulated across different sample types, indicating a common redox stress response. Nonetheless, the alteration of relevant metabolites was not consistent across different sample types, showing differential mechanisms of NGLY1-deficiency-induced stress responses. Differential alternations of glutathione metabolism were also reported in transcriptomics studies across different types of patient-derived cell lines^135^.

We also found that the iPSC proteome did not demonstrate significant alteration in immune relevant pathways whereas the NPC proteome demonstrated upregulation of interleukin signaling, suggesting potential immune activation induced by dysregulated nerve function. The activation of the ubiquitination signaling pathway and NRF2-mediated antioxidant response were molecular phenotypes of the CP1 patient’s NPCs. The upregulation of diverse ubiquitination pathways could be a consequence of disrupted deglycosylation pathways, as ubiquitination is a prerequisite for NGLY1-catalyzed deglycosylation. NRF2-mediated oxidative responses were also a consequence of NGLY1 dysfunction. Previous studies have suggested that deglycosylation is a crucial step for the maturation of the NRF family. Nonetheless, as the relevant pathways were not significantly alternated in patient iPSCs, this may be a downstream phenotype.

NGLY1 deficiency is a rare disease with extensive and diverse symptoms that has suffered from limited therapeutic development and a lack of common biomarkers. Our results show that patients’ plasma exhibit individualized molecular phenotypes. The clustering of both plasma metabolome and proteome fail to separate patients and healthy completely (Supplementary Fig. 3A, 4A, 5A, 7A). The cytokine profiles also demonstrate highly personalized immunophenotypes (Supplementary Fig. 10A). Potential reasons for such diversity could be the differential downstream pathways regulated by NGLY1 in individual patients. The patient-derived cell models combined with gene-editing technology provided a direct solution to such problems, enabling study of the functional consequences of individual patient heterogeneity. As more patients are included in the generation of personalized cellular models, additional personalized pathway responses can be discovered and targeted for treatment.

## Conclusion

This study reports the most comprehensive and updated multi-omics study of NGLY1 deficiency to date in a cohort that covers a majority of reported cases worldwide. CP1 patient-derived cellular models enabled much deeper characterization of dysregulated pathways in NGLY1 deficiency. The differential metabolites and proteins identified in both plasma samples and cellular models revealed the alteration of diverse pathways in NGLY1 deficiency, including lipid metabolism, amino acid metabolism, biogenic amine synthesis, glutathione metabolism, and immune response. The plasma metabolome and proteome demonstrated significant individualization of patients’ molecular phenotypes. The personalized iPSC/NPC metabolome and proteome data revealed that the onset of immune response may be a consequence of other dysregulated pathways. These findings provide useful insight into the global proteomic and metabolic alterations induced by NGLY1 deficiency and will facilitate therapeutic and diagnosis development for the disease. This approach is also transferable to the study of other underdiagnosed rare diseases and can be used to shed light on the pathologic and therapeutic investigation of other conditions.

## Materials and Methods

### Chemical materials

MS-grade water (7732-18-5), methanol (A456-500), acetonitrile (A9554), and acetone (67-64-1) were purchased from Fischer Scientific (Morris Plains, NJ, USA). MS-grade acetic acid (64-19-7) was purchased from Sigma Aldrich (St. Louis, MO, USA).

### Metabolomics analysis

We profiled the plasma samples using both targeted and untargeted approaches. For targeted profiling, we used the “AbsoluteIDQ p400 HR assay” kit to quantify 396 plasma metabolites including sugars, amino acids and biogenic amine derivatives and lipids. In parallelly, we used the Lipidyzer platform (SCIEX) to quantify 723 lipid species across 13 lipid classes. The untargeted LC-MS approach provided a broader coverage of the plasma metabolome. We also used the untargeted approach to profile plasma metabolome and the personalized disease modeling cellular metabolome to validate the comprehensive molecular phenotypes of plasma and trace the progression of the disease in patient-derived cell models.

### Targeted metabolomics

10 μL plasma were prepared and analyzed following the sample preparation procedures and MS/MS (Tandem Mass Spectrometry)-based analytical methods provided with “AbsoluteIDQ® p400 HR assay” kit. LC-MS and flow injection analysis (FIA) data were acquired on a Q Exactive HF mass spectrometer (Thermo Scientific). The raw data was processed by Xcarlibur (Thermo Scientific) and MetIDQ (Biocrates). The complete list of quantified plasma metabolites was reported in Supplementary Table 2.

### Lipidomics profiling of plasma samples

Lipids were extracted and analyzed using a standard protocol^135^. A mixture of methyl tertiary-butyl ether, methanol and water was used to extract lipids from 40□µL of plasma by biphasic separation. Lipids were then analyzed with the Lipidyzer platform consisting in a DMS device (SelexION Technology, SCIEX) and a QTRAP 5500 (SCIEX). Lipids were semi-quantified using a mixture of 58 labeled internal standards provided with the platform. The quantified plasma lipids were reported in Supplementary Table 4.

### Untargeted metabolomics

200 μL plasma was treated with 800 μL 1:1:1 (v/v) acetone:acetonitrile:methanol solvent mixture, spiked-in with 15 analytical-grade internal standards, mixed for 15 min at 4°C, and incubated at −20°C for 2 h to allow for protein precipitation. iPSCs/NPCs were washed 3 times with PBS, then lysed by 90% methanol, spiked-in with 15 analytical-grade internal standards, vortexed for 10 s, sonicated for 30 s, spin down and cooled down on ice for 30 s, repeat 3 times. The supernatant of plasma/cell lysate was collected after centrifugation at 10,000 rpm for 10 min at 4°C and evaporated under nitrogen to dryness. The dry extracts were reconstituted with 100 μL 1:1 methanol:water before analysis. A quality control (QC) sample was generated by pooling all the plasma samples injected between every 10 sample injections to monitor the consistency of the retention time and the signal intensity. The QC sample was also diluted two, four, and eight times to determine the linear-dilution effect of metabolic features. Untargeted metabolomics was performed with an optimized pipeline on a broad-spectrum LC–MS platform^137^. In brief, complementary reverse-phase liquid chromatography (RPLC) and hydrophilic interaction liquid chromatography (HILIC) separations were applied in parallel to maximize metabolome coverage. Data were acquired on a Q Exactive plus mass spectrometer (Thermo Scientific). The instrument was operated in full mass-spectroscopy scan mode. Tandem mass spectroscopy data was acquired at various collision energies on pooled quality control samples. LC–MS data were processed using Progenesis QI (Nonlinear Dynamics) and metabolic features were annotated by matching retention time and fragmentation spectra to authentic standards or to public repositories by metID pipeline available in https://jaspershen.github.io/metID/ and validated manually^129^. The untargeted metabolomics profiles of plasma and iPSCs/NPCs were reported in Supplementary Table 6, 8.

### Proteomics analysis

LC-MS based Tandem Mass Tag (TMT) labeled identification and quantification of proteome were applied for the proteomics profiling of both plasma samples and disease-modeling cell lysates. In brief, for patient-derived cells, cell pellets were lysed in a fresh prepared cell lysis buffer containing 6 M Gdmcl, 10 mM TCEP, 40 mM CAA, 500 mM Tris (pH 8.5). For plasma proteome, 25 μL plasma were depleted using an Agilent Mars human 14 column (4.6 mm x 50mm). The unbound fraction was collected for buffer exchange with the fresh cell lysis buffer for further proteome analysis as previously described^130^. After protein reduction and alkylation, protein concentration was measured using the BCA kit (ThermoFisher). 100 mg protein pellet was used for trypsin digestion (1:50) overnight at 37 °C. Peptides were cleaned up using Waters HLB column. Samples were randomized and labeled using TMT10 Plex (ThermoFisher) in 100 mM TEAB buffer. An equal amount of peptides of each sample were pooled together as a reference. Data acquisition was operated on a NanoAquity 2D nanoLC (Waters) directly coupled in-line with a linear trap quadrupole (LTQ)-Orbitrap Velos instrument (Thermo Scientific) via Thermo nanoelectrospray source. 15 μg multiplexed sample was loaded on 2D-LC for online fractionation. Peptides were separated by reverse-phase chromatography at high pH in the first dimension, followed by an orthogonal separation at low pH in the second dimension. The raw data were processed with the Proteome Discoverer 2.1 (Thermo), using UniProtKB/Swiss-Prot database, applying 1% FDR. Due the limited sample amount, plasma sample NGLY1-60 was excluded for proteomics profiling. The plasma proteomics profiles were reported in Supplementary Table 9. The proteomics profiles of iPSCs/NPCs were reported in Supplementary Table 12, 13.

### Immune protein measurements

The 62 plex-Luminex antibody-conjugated bead capture assay (Affymetrix) was used to characterize blood levels of immune proteins. The assay was performed by the Stanford Human Immune Monitoring Center. The protocol is available in https://iti.stanford.edu/himc/protocols.html. The cytokine profiles were reported in Supplementary Table 11.

### Data analysis and integration

The statistical analysis and data visualization was performed using R language (version 3.6.0) and R studio (version 1.2.5019). The directory used R packages included plyr, stringr, dplyr, purrr, readr, readxl, tidyr, tibble, ggplot2, ggraph, igraph, tidygraph, pheatmap, ComplexHeatmap, ggrepel, EnhancedVolcano, magrittr.

A linear mixed model was applied to plasma metabolome and proteome data to adjust potential bias induced by the unmatching age distribution between patients and controls (Supplementary Fig. 2B, Table 1). All t-tests were two-tailed, with a P-value of less than 0.05 considered significant. The altered metabolites and proteins were used for pathway enrichment analysis, correlation analysis and network analysis with IPA (QIAGEN Inc., https://www.qiagenbioinformatics.com/products/ingenuitypathway-analysis)140. Tissue enrichment was conducted by mapping the NGLY1-relavant proteins with tissue specific quantitative proteome^130^.

### Generation of iPSCs and NPCs

Patient’s skin biopsy samples were collected in Stanford Hospital and fibroblast cells were derived in Stanford Cytogenetics Laboratory. The iPSCs were generated with the Sendaivirus delivery system^138^ using the CytoTune-iPS 2.0 Sendai Reprogramming Kit (ThermoFisher Scientific). Briefly, the fibroblast cells were maintained in DMEM containing 10% fetal bovine serum. During the reprogramming, 2×10^5^ cells were transduced with Sendai virus with MOI of 5:5:3 (KOS:c-Myc:Klf4) as instructed in the manual. The cells were maintained in fibroblast media for 6 days then passaged onto Matrigel-coated dishes for further reprogramming in Essential 6 medium plus human bFGF (ThermoFisher Scientific). The iPSC colonies were manually picked onto Matrigel-coated plates to generate stable iPSC lines. The iPSCs were maintained in mTeSR1 medium (Stemcell Technologies) and routinely passaged every 4∼6 days with Versene (EDTA) solution (Lonza). Selected iPSC clones were further validated for expression of pluripotency markers with immunostaining (Supplementary Fig. 2) and NGLY1 mutations were confirmed with Sanger sequencing. The karyotyping of the iPSCs were checked by G-banding analysis in WiCell institute.

### Ngly1 mutation correction with CRISPR-Cas9 gene editing in iPSCs

For Ngly1 gene editing, the guide RNA sequence was cloned into pX459 vector for either exon 8 (c.1205_1207delGAA) or exon 11 (c.1624C>T) mutation correction. Patient-derived iPSCs harboring Ngly1 mutations were dissociated with Accutase into single cells, and 2×10^6^ iPSCs were mixed with pX459-sgRNA vector and ssDNA repair donor template and electroporated with Lonza 4D-Nucleofector system using the P3 Primary Cell Nucleofector Kit (Lonza). The iPSCs were then recovered in mTeSR1 medium for 48 hours before adding puromycin for 3-day selection. The survived iPSCs were dissociated with Accutase and plated at a very low density for subcloning. Each individual iPSC colony was expanded and screened with Sanger sequencing for mutation correction.

### Feeder-free differentiation of iPSCs to NPCs

The feeder-free NPC differentiation was achieved by dual-SMAD signaling inhibition modified from the publication from Dr. Studer lab^139^. iPSC monolayer cultures were incubated in KO-DMEM medium containing 20% knockout serum replacement (KSR, ThermoFisher) with 10uM SB431542, 100nM LDN-193189, Glutamax (1/100), NEAA (1/100) and beta-mercaptoethanol (0.1 mM). The cells were fed with fresh media every other day and collected seven days after differentiation.

## Acknowledgements

We thank the following people for their essential contributions to this paper: Jing Liang and Yael Rosenberg-Hasson at Stanford Human Immune Monitoring Center (HIMC) for cytokine quantification; Lulu Cao and Christine Yeh for the pilot test of iPSC proteomics analysis; Alexander Honkala, Dan Li, Yu Zhang, Liang Liang and Ashley Tan for reviewing the manuscripts; Ashley Tan for validating data analysis process. This work was supported by Grace Science Foundation.

## Supplementary figures

**Fig S1.**
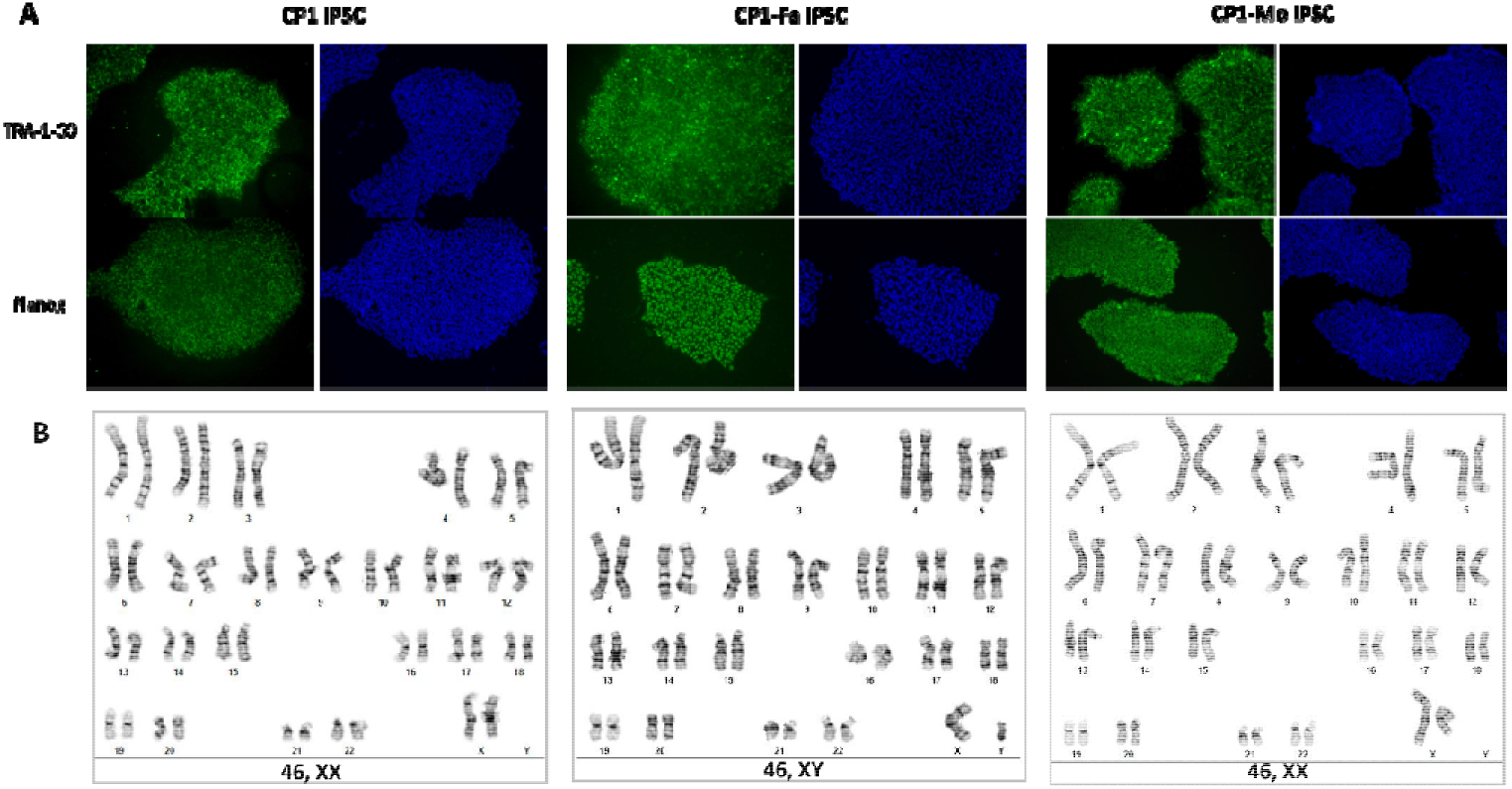
Characterization of patient-derived iPSCs. A) iPSCs from patient CP1’s family (CP1, CP1-Fa, CP1-Mo) were confirmed with pluripotency markers expression (TRA-1-60, Nanog). B) iPSCs from CP1’s family were confirmed with normal karyotyping (by G-banding).

**Fig S2.**
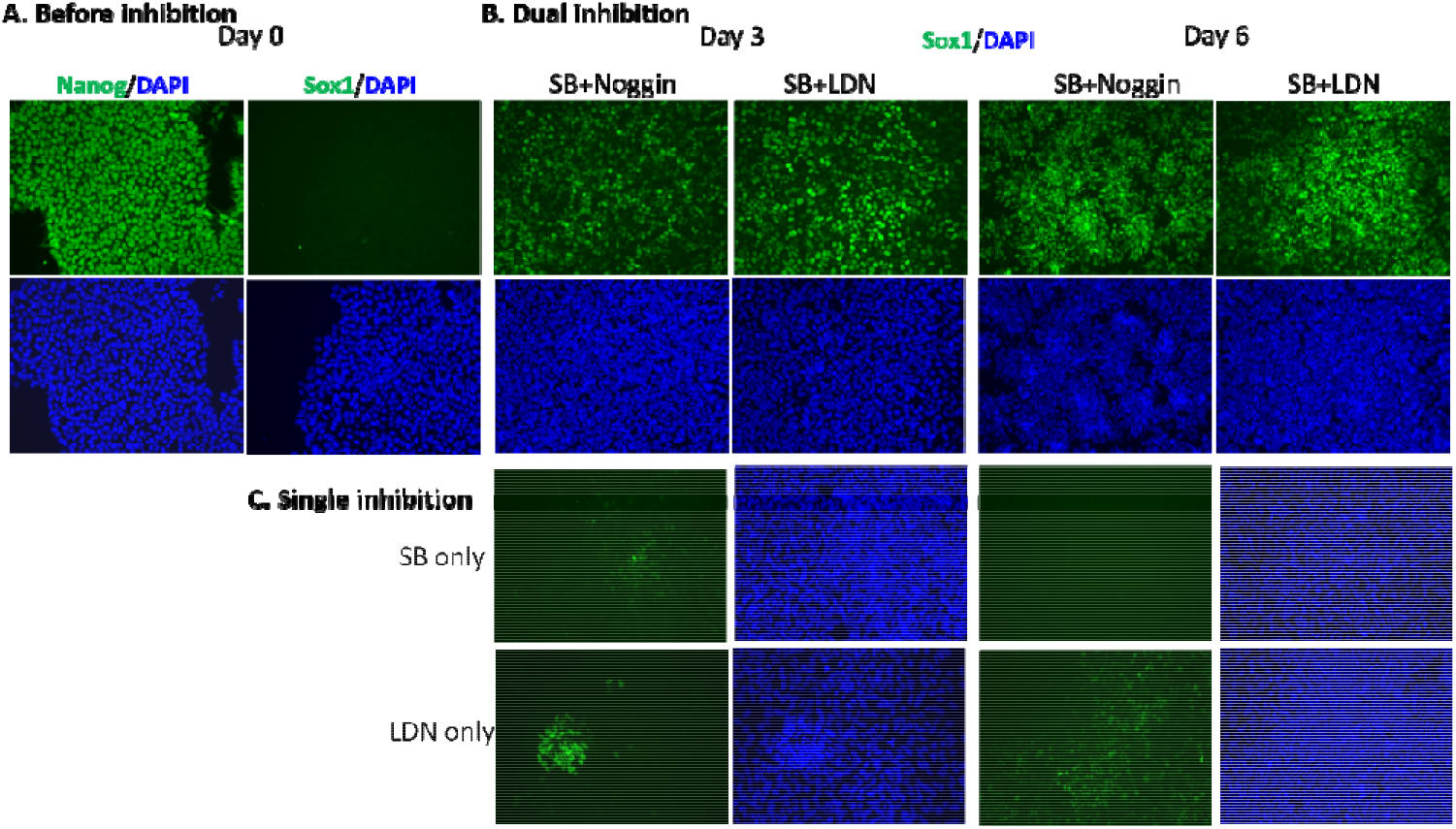
The induction of iPSCs by dual and single inhibition. A) Fluorescence imaging of iPSCs before inhibition on day 0. B) The induction of iPSCs to NPCs by dual inhibition (SB+Noggin, SB+LDN). C) The induction of iPSCs to NPCs by single inhibition (SB, LDN).

**Fig S3.**
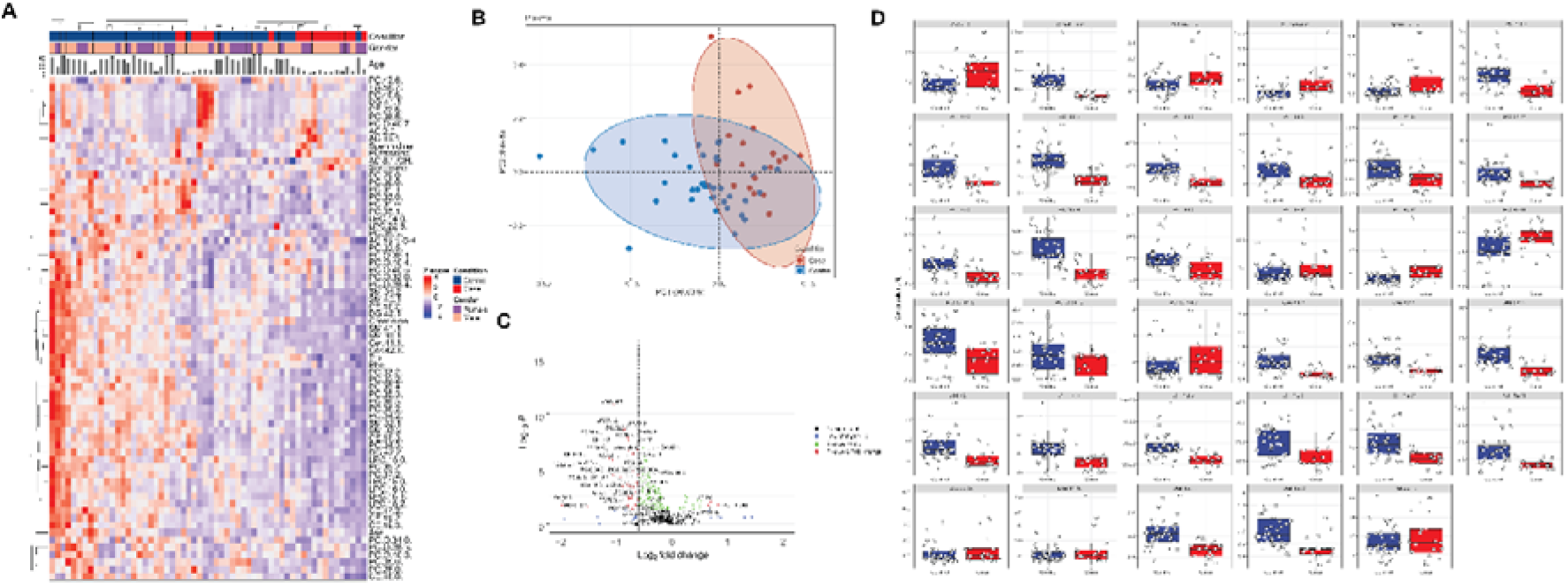
Summary of the targeted metabolomics profiling of NGLY1 deficiency plasma samples. A) The hierarchical clustering of metabolites with significant variation (P < 0.05) between patients and controls. B) PCA of targeted metabolomics profiles demonstrated partial separation of patients and controls. C) Volcano plot of plasma metabolites with significant variation and fold changes between patients and control. D) Boxplots of plasma metabolites with highest variation between patients and controls.

**Fig S4.**
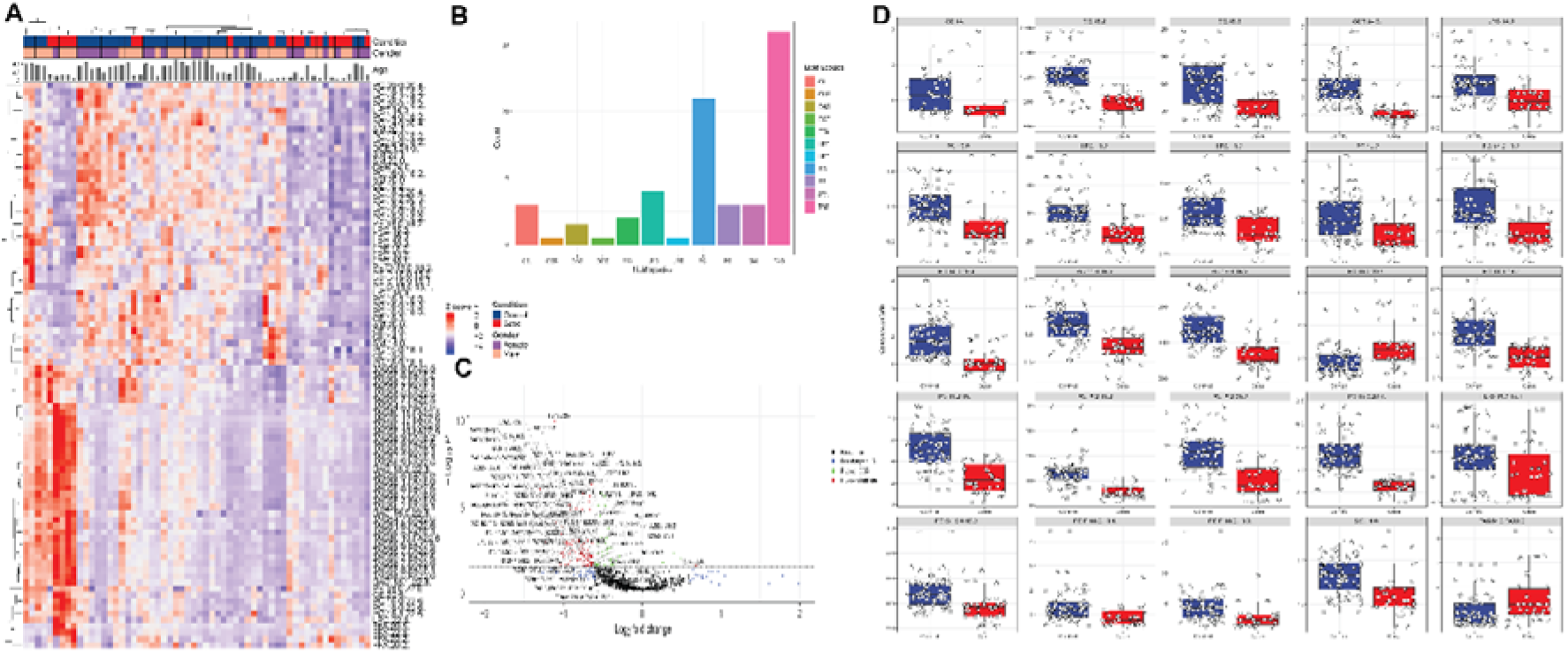
Summary of the lipidomics profiling of NGLY1 deficiency plasma samples. A) The hierarchical clustering of lipids with significant variation (P < 0.05) between patients and controls. B) Amounts of lipid species with significant variation between patients and controls (P < 0.05). C) Volcano plot of plasma lipidome. The labeled lipids show a fold change > 1.5 between patients and control (P < 0.05). D) Boxplots of plasma lipids with fold change > 1.5 between patients and controls (P < 0.05).

**Fig S5.**
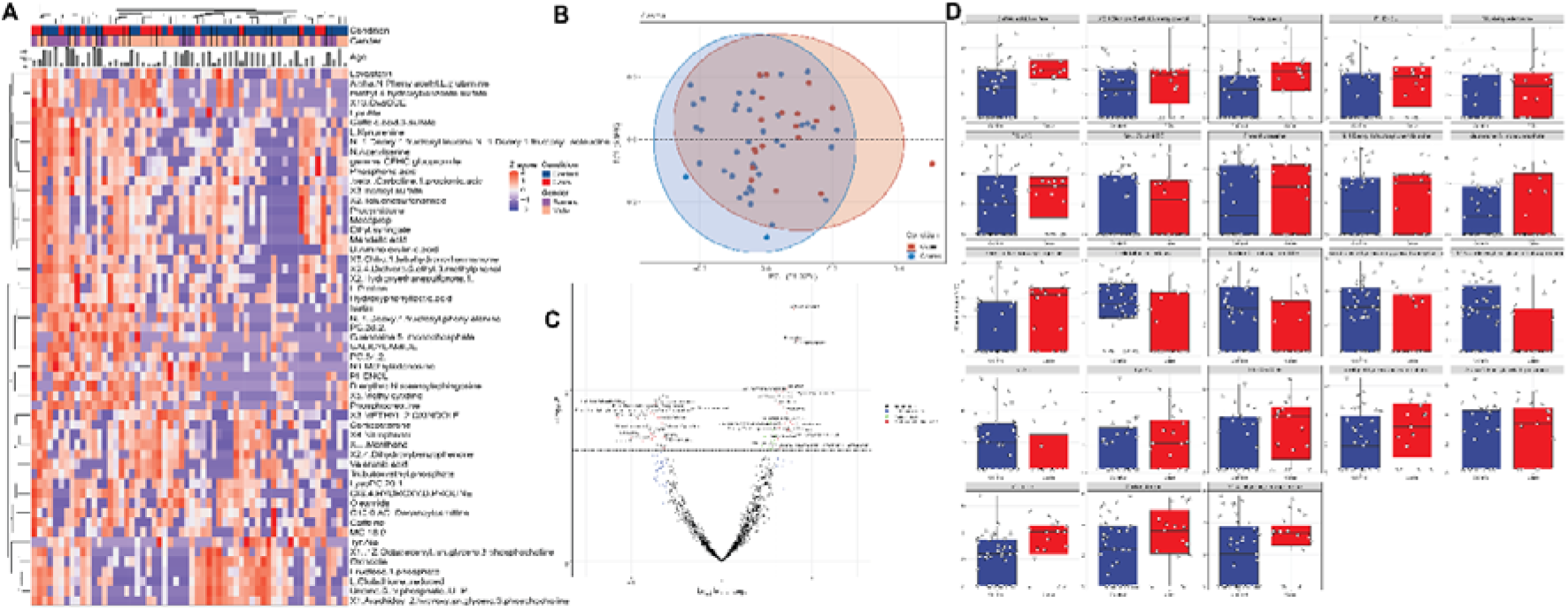
Summary of the untargeted metabolomics profiling of NGLY1 deficiency plasma samples. A) The hierarchical clustering of metabolites with significant variation (P < 0.05) between patients and controls. B) PCA of untargeted metabolomics profiles demonstrated minor separation of patients and controls. C) Volcano plot of plasma metabolome. The labeled lipids show a fold change > 1.5 between patients and control (P < 0.05). D) Boxplots of plasma metabolites with most significant variation between patients and controls.

**Fig S6.**
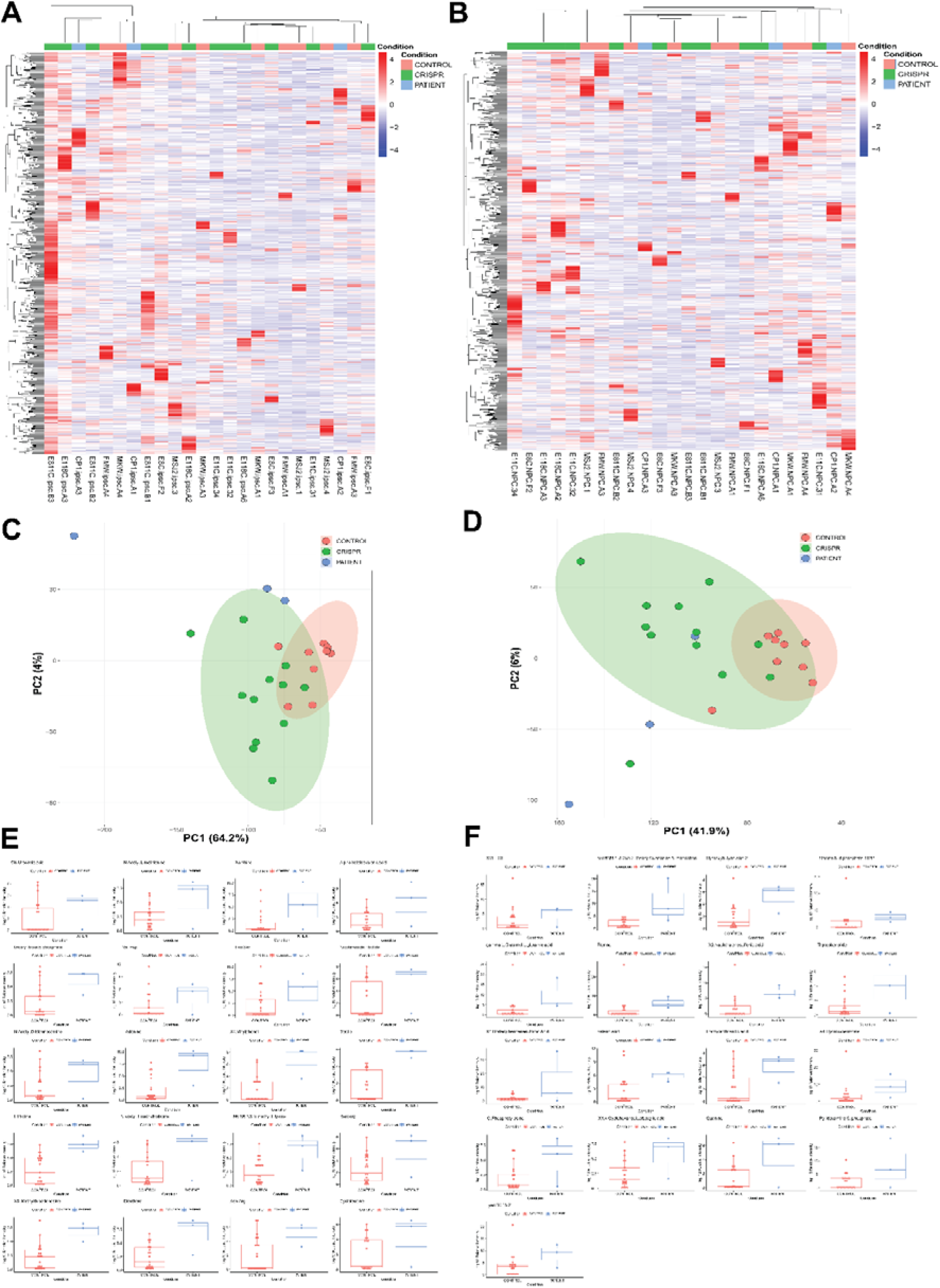
Summary of the untargeted metabolomics profiling of personalized disease-modeling iPSC and NPC. A) The hierarchical clustering of iPSC metabolome. B) The hierarchical clustering of NPC metabolome. C) PCA of iPSC metabolome demonstrated minor separation between patients, isogenic controls, and external controls. D) PCA of NPC metabolome demonstrated minor separation between patients, isogenic controls, and external controls. E) Boxplot of iPSC metabolites with vast fold changes between patients and control. F) Boxplot of NPC metabolites with vast fold changes between patients and control.

**Fig S7.**
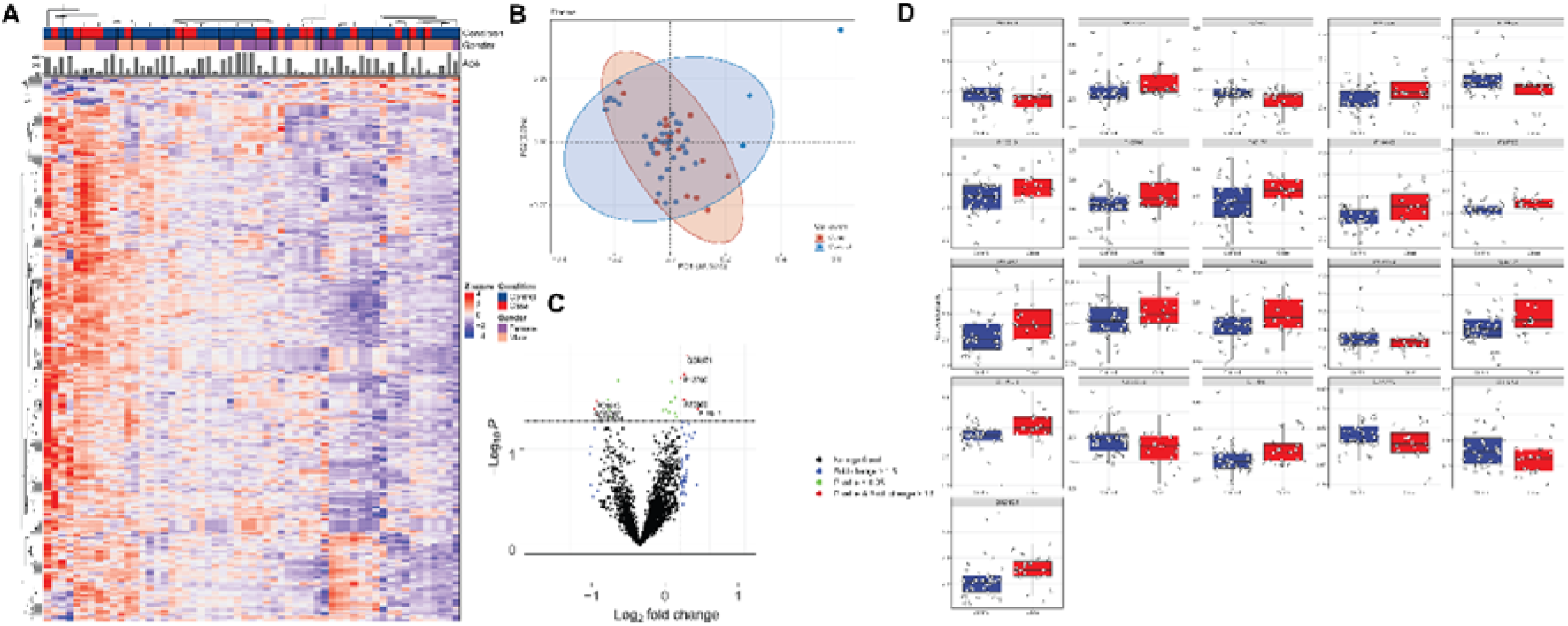
Summary of proteomics profiling of NGLY1 deficiency plasma samples. A) The heatmap of plasma proteome. B) PCA of proteomics profiles demonstrated minor separation between patients and controls. C) Volcano plot of plasma proteins. The labeled cytokines show a fold change > 1.5 between patients and control (P < 0.05). D) Boxplots of plasma proteins with most significant fold changes and variation between patients and controls.

**Fig S8.**
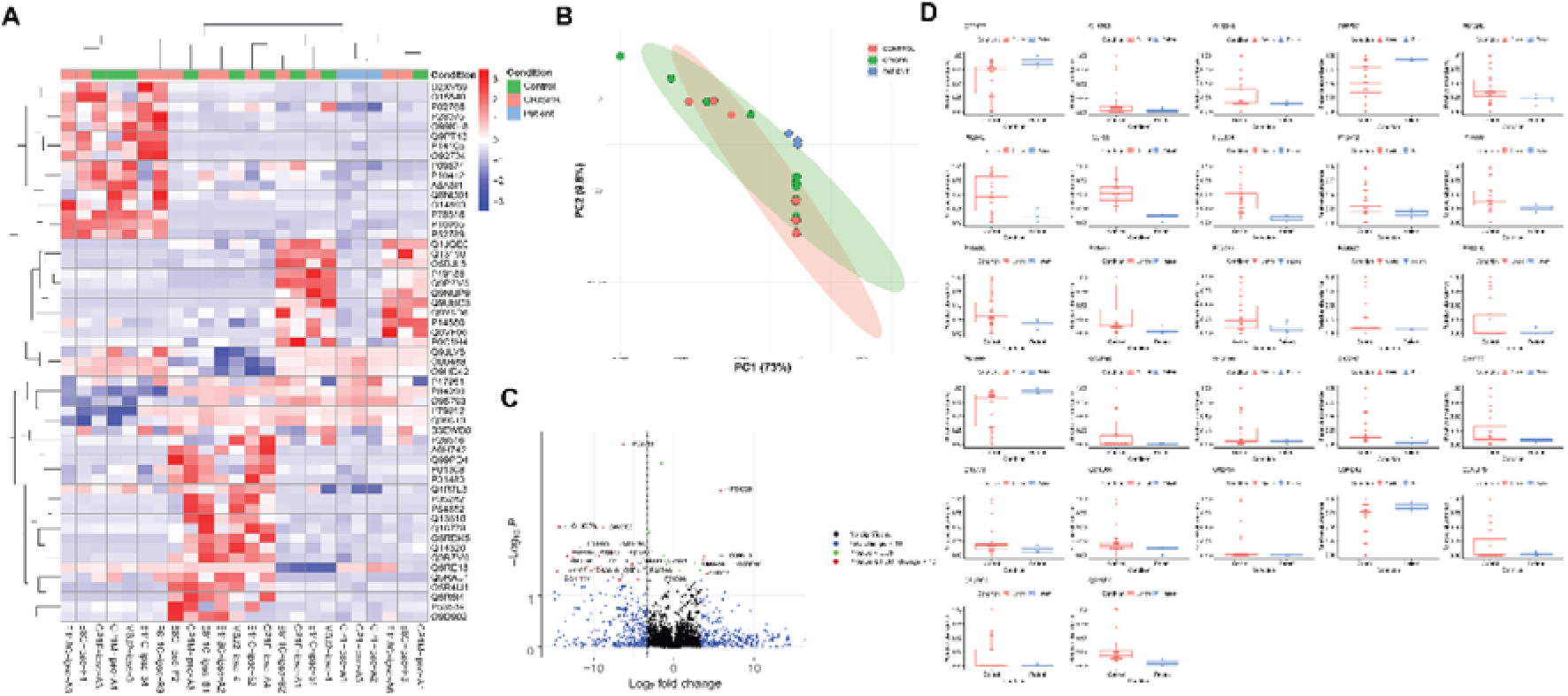
Summary of proteomics profiling of disease-modeling iPSC derived from Patient CP1’s family and external controls. A) The hierarchical clustering of proteins with significant variation (P < 0.05) between patients and controls. B) PCA of iPSC proteomics profiles demonstrated partial separation between patients and controls. C) Volcano plot of iPSC proteins with significant variation and fold changes between patients and control. D) Boxplots of iPSC proteins with most significant fold changes and variation between patients and controls.

**Fig S9.**
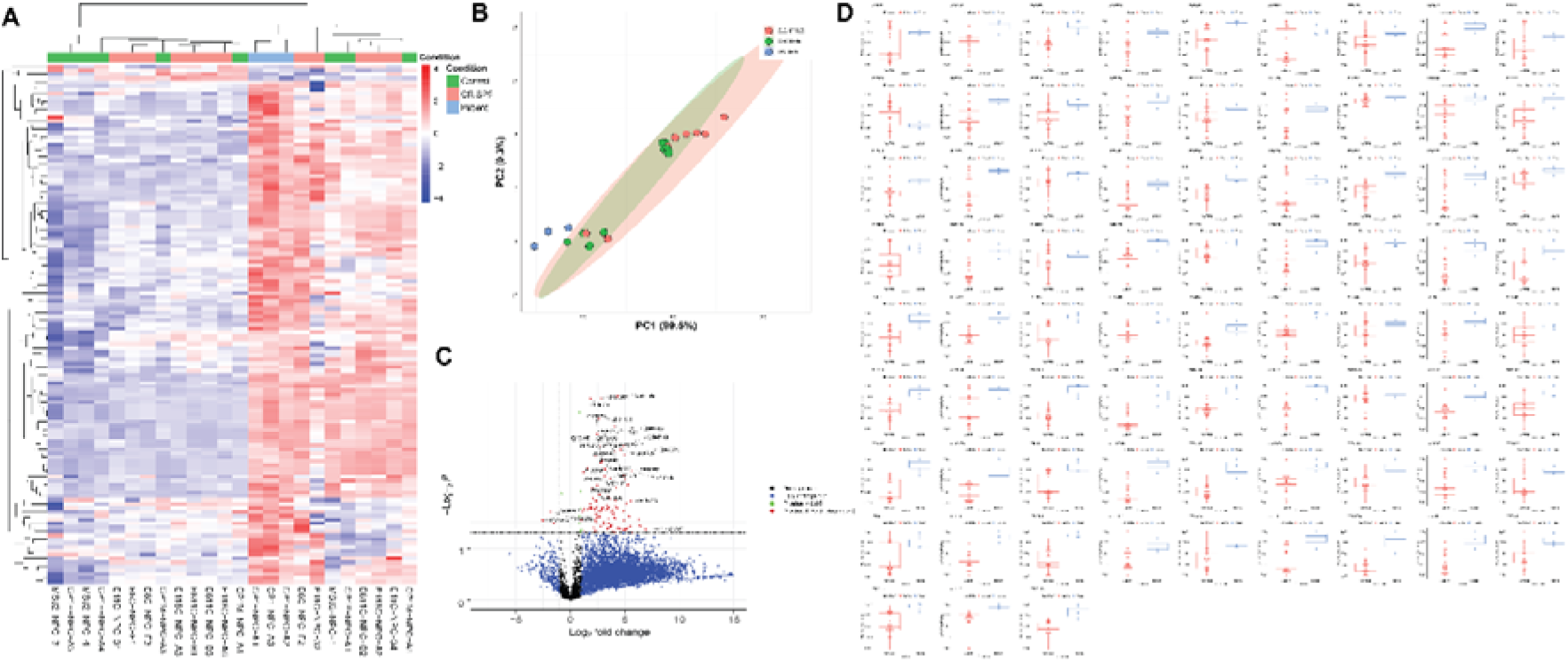
Summary of proteomics profiling of disease-modeling NPC derived from Patient CP1’s family and external controls. A) The hierarchical clustering of proteins with significant variation (P < 0.05) between patients and controls. B) PCA of NPC proteomics profiles demonstrated partial separation between patients and controls. C) Volcano plot of NPC proteins with significant variation and fold changes between patients and control. D) Boxplots of NPC proteins with most significant fold changes and variation between patients and controls.

**Fig S10.**
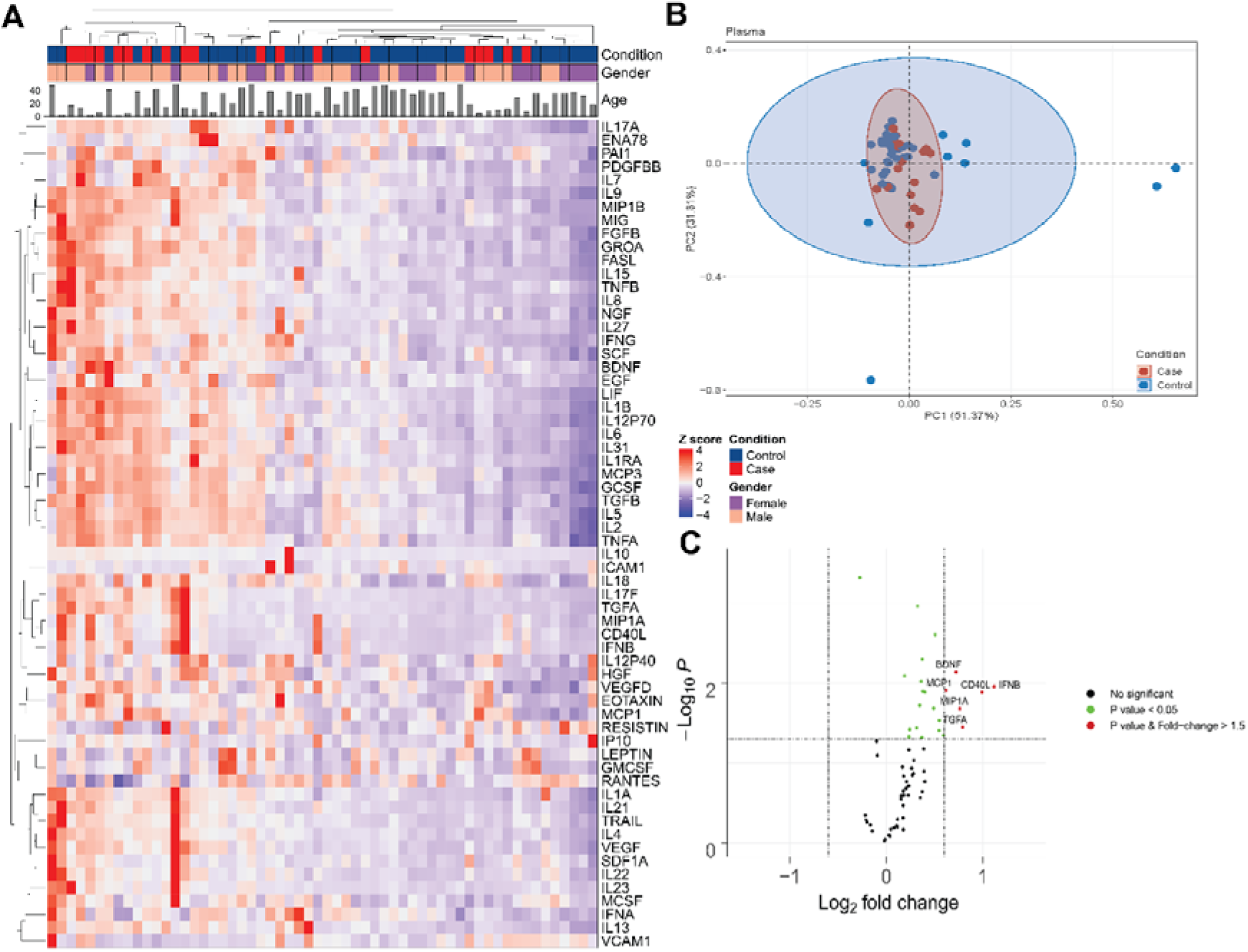
Summary of cytokine measurement of NGLY1 biobank plasma samples. A) The heatmap of cytokines. The hierarchical clustering shows partial separation between patients and controls. B) PCA of cytokine profiles demonstrated minor separation between patients and controls. C) Volcano plot of cytokine profiles. The labeled cytokines show a fold change > 1.5 between patients and control (P < 0.05).

## Notes

**Ethics declarations:** Competing interests The authors declare no competing interests.

### Competing Interest Statement

The authors have declared no competing interest.

